# Structural elucidation of the heterodimeric *cis*-prenyltransferase NgBR/DHDDS complex reveals novel insights in regulation of protein glycosylation

**DOI:** 10.1101/2020.06.03.132209

**Authors:** Ban Edani, Kariona A. Grabińska, Rong Zhang, Eon Joo Park, Benjamin Siciliano, Liliana Surmacz, Ya Ha, William C. Sessa

## Abstract

*Cis*-prenyltransferase (*cis*-PTase) catalyzes the rate-limiting step in the synthesis of glycosyl carrier lipids required for protein glycosylation in the lumen of endoplasmic reticulum. Here we report the crystal structure of the human NgBR/DHDDS complex, which represents the first atomic resolution structure for any heterodimeric *cis*-PTase. The crystal structure sheds light on how NgBR stabilizes DHDDS through dimerization, participates in the enzyme’s active site through its C-terminal -RXG- motif, and how phospholipids markedly stimulate *cis*-PTase activity. Comparison of NgBR/DHDDS with homodimeric *cis*-PTase structures leads to a model where the elongating isoprene chain extends beyond the enzyme’s active site tunnel, and an insert within the α3 helix helps to stabilize this energetically unfavorable state to enable long chain synthesis to occur. These data provide unique insights into how heterodimeric *cis*-PTases have evolved from their ancestral, homodimeric forms to fulfill their function in long chain polyprenol synthesis.

## Introduction

Most members of the highly conserved *cis*-prenyltransferase (*cis*-PTase) family catalyze the sequential condensation of an allylic pyrophosphate with a variable number of 5-carbon isopentenyl pyrophosphates (IPPs), resulting in the formation of polyprenol pyrophosphates with *cis*-double bonds ^1–4^. In eukaryotes, the endoplasmic reticulum (ER) associated *cis*-PTase is the first enzyme committed to the synthesis of dolichol phosphate (DolP) ^4–8^, an indispensable lipid carrier for protein N-glycosylation, O-mannosylation, C-mannosylation and GPI-anchor formation ^9,10^. In contrast to mammalian *cis*-PTase, undecaprenyl pyrophosphate synthase (UPPS), a bacterial *cis*-PTase, is essential for cell wall synthesis ^11^. Eukaryotic *cis*-PTases have the extraordinary ability to synthesize long-chain (14-24 C_5_ isoprene units) or very-long-chain (>2,000 C_5_) products ^12^, whereas bacterial, some protistic, archaeal and plant enzymes mainly produce medium-chain (9-11 C_5_), or short-chain (2-5 C_5_), products ^2,3,10,13–16^.

Although the structure and mechanism of homodimeric, bacterial *cis*-PTases, have been extensively studied ^1,17–22^, the eukaryotic enzyme’s unique mechanism has remained elusive until recently. Dehydrodolichyl diphosphate synthase (DHDDS) or its yeast orthologues Rer2 and Srt1, retain most of the active site residues in common with bacterial *cis*-PTase ^4–6^, however, as purified, have very little enzymatic activity ^23^. The full activity of *cis*-PTase requires the interaction of DHHDS with an additional subunit, originally identified as NgBR (Nogo-B receptor) or Nus1 in yeast ^24–31^. Within eukaryotic cells, DHDDS is stabilized by the association with NgBR ^24^. Part of NgBR shares sequence and structural homology with bacterial *cis*-PTase ^30,32,33^, but lacks most catalytic residues. Outside the *cis*-PTase homology domain, NgBR has a N-terminal membrane-binding region ^24^ (Fig. 1A). Various roles have been proposed for NgBR and NgBR-like proteins in the membrane association and function of eukaryotic *cis*-PTases and a subgroup of archaeal enzymes that share this heteromeric arrangement ^24,33–37^. We have previously shown that NgBR and Nus1 are is indispensable for *cis*-PTase activity in humans and in yeast, respectively ^25,33^.

**Figure 1.**
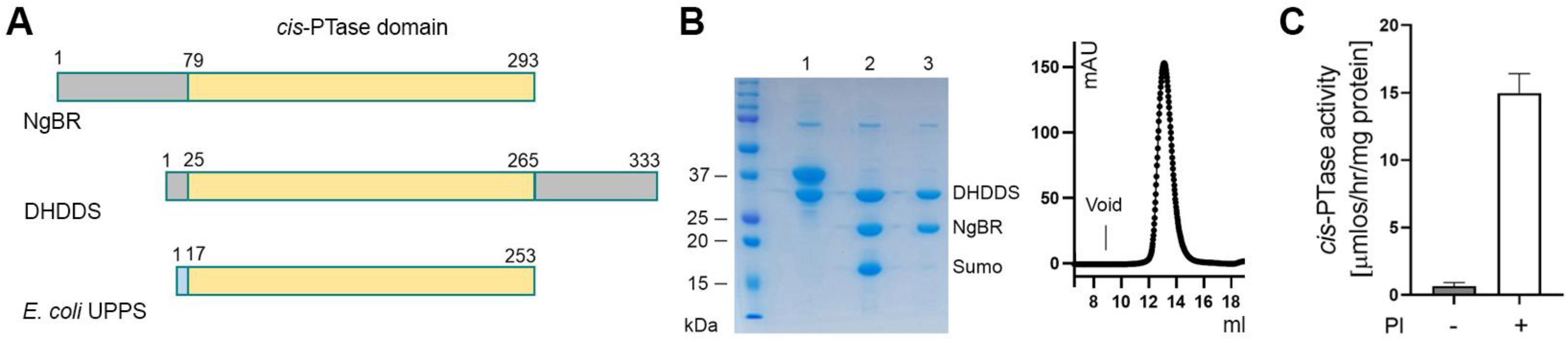
The catalytic core domain of human *cis*-PTase. (A) Comparing the domain structure of NgBR, DHDDS and *E.coli* UPPS. The *cis*-PTase domain is colored yellow; N- and C-terminus of NgBR and DHDDS, gray; N-terminus of *E.coli* UPPS, blue. (B) Purification of NgBR/DHDDS complex. Left Panel: Coomassie-stained SDS/PAGE showing purification steps. Lane 1: uncleaved 6HIS-SUMO-NgBR/DHDDS complex, Lane 2: cleaved NgBR/DHDDS complex and SUMO, Lane 3: NgBR/DHDDS complex after removing SUMO. Right panel, Size exclusion chromatography profile of the purified complex after cleavage with SUMO protease. (C) Stimulation of *cis*-PTase activity of NgBR/DHDDS complex by phosphatidylinositol (PI). The values are means ±S.D. of eight independent measurements from two independent purifications.

Exome sequencing of patients with congenital disorders of glycosylation (CDG) has identified a number of disease-causing mutations in both DHDDS and NgBR ^25,38–42^. A homozygous mutation in NgBR at Arg-290 (NgBR R290H mutation), occurs in a conserved C-terminal -RXG- motif that is shared amongst NgBR orthologs as well as in bacterial homodimeric *cis*-PTases, but is absent in DHDDS and its orthologs, suggesting that the C tail of NgBR participates in *cis*-PTase function. Biochemical characterization of the R290H mutant revealed this mutation impairs IPP binding and markedly reduces catalytic activity but does not influence its interaction with DHDDS ^25,33^ raising the possibility that NgBR could directly contribute to substrate binding and catalysis through its -RXG- motif ^21,43,44^. Another recent exome sequencing study identified multiple mutations in NgBR that appear to cause Parkinson’s disease ^45^, although it is uncertain whether these mutations affect *cis*-PTase activity.

Here, we report the first crystal structure of a heteromeric, human *cis*-PTase NgBR/DHDDS complex solved at 2.5 Å resolution. Besides proving a catalytic role of NgBR’s -RXG- motif, structural-functional analyses have unveiled several unique attributes of the complex that were not predicted based on the structures of UPSS. This includes a unique C-terminal clamp in DHDDS that contributes to heterodimerization; the mechanistic basis for a Parkinson’s disease causing, loss of function mutation; a novel N-terminal membrane sensor critical for lipid activation of *cis*-PTase activity and a critical structural feature in DHDDS that impacts product polyprenol chain length. Thus, our crystal structure provides unique insights into how heterodimeric mammalian *cis*-PTases have evolved from their ancestral, homodimeric forms to synthesize long chain polyprenols and dolichol, lipids essential for protein glycosylation reactions.

## Results and Discussion

### The core catalytic domain of NgBR/DHDDS complex

The rate-limiting step in dolichol biosynthesis in the ER is catalyzed by *cis*-PTase, which consists of NgBR and DHDDS subunits ^24,25^. Unlike homodimeric *cis*-PTase found in bacteria that synthesizes a medium-chain-length polyprenol pyrophosphate (e.g., *E. coli* UPPS; 11 isoprene units), human *cis*-PTase preferentially catalyzes the condensation of sixteen IPP molecules with a single farnesyl pyrophosphate (FPP), generating a long-chain reaction product (i.e. 19 isoprene units) ^10,11^. To facilitate structural and mechanistic characterization of the heteromeric *cis*-PTase, we co-expressed a polyHis- and SUMO-tagged human NgBR ^46^, which has the N-terminal 78 amino acids deleted (Fig. 1A), with full-length human DHDDS in *E. coli*, and purified the protein complex to homogeneity in milligram quantities (Fig. 1B). Consistent with the model where NgBR’s N-terminal region mainly functions as a membrane anchor ^24^, the N terminal truncated NgBR/DHDDS complex is as active enzymatically as the full-length protein complex purified from mammalian cells ^33^. Also like the full-length protein, the activity of the truncated NgBR/DHDDS was potently stimulated by phosphatidylinositol (PI), a phospholipid abundantly present in the ER membrane (Fig. 1C) ^33^. Reverse phase thin layer chromatography (RPTLC) confirmed that NgBR N-terminal truncation did not alter the range, or relative abundance, of various long-chain polyprenols generated from the *in vitro* reaction (see below).

### The heterodimeric structure

The core catalytic domain of the NgBR/DHDDS complex was crystallized in the presence of IPP and Mg^2+^. The crystal belongs to space group R32, and the asymmetric unit contains a single NgBR^79-293^/DHDDS heterodimer (hereinafter referred to as NgBR/DHDDS). The structure was determined by molecular replacement using *E.coli* UPPS (PDB entry 1X06) and *S.cerevisiae* Nus1 (PDB entry (6JCN) as search probes for DHDDS and NgBR, respectively (Fig. 2A; supplementary Table 1) ^32,47^. Clear electron densities enabled us to model several functionally important elements that were absent in the search probes (supplementary Fig. 1–3). In NgBR, these structural elements include an N-terminal α-helix (α1, residues 82-93) and an extended C-terminal segment (residues 286-293) that contains the catalytically essential -RXG- motif. Furthermore, the two outermost β-strands (βC, βC’) observed in Nus1 are disordered (residues 230-244; supplementary Fig. 3). The peptide segment following the β-strands forms a new helix (α4, residues 179-186) and partially covers the hydrophobic cavity previously proposed to constitute the binding site for farnesylated Ras ^48^. In DHDDS, the additional structures include a conserved N-terminal segment (residues 1-24) that reaches into the hydrophobic tunnel of the active site, and a pair of long α-helices (residues 251-333) toward the C-terminus that wrap around the protein complex.

**Figure 2.**
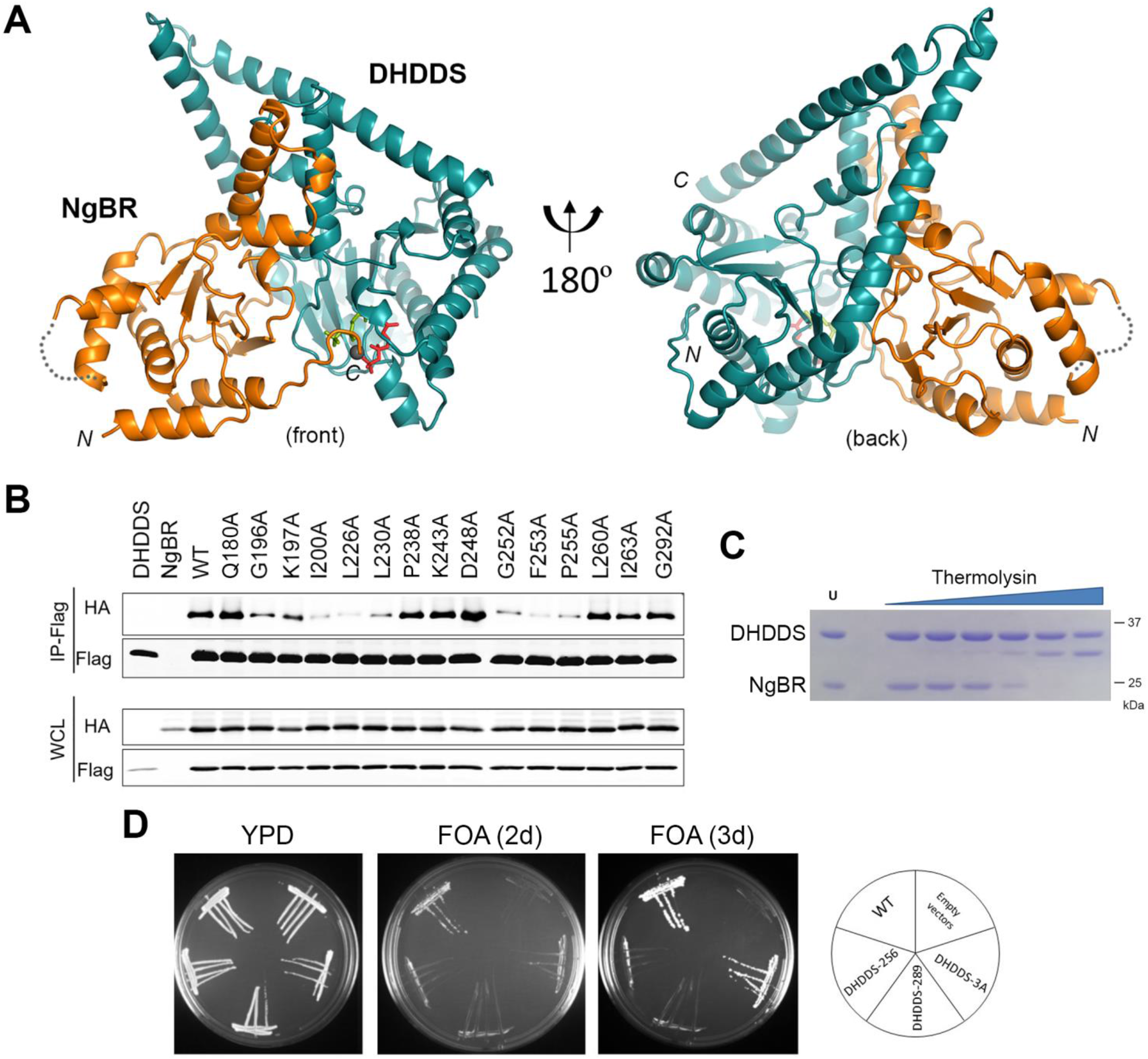
The overall structure of NgBR/DHDDS heterodimer. (A) Ribbon diagrams showing the front and back of the heteromeric complex. NgBR is colored in orange and DHDDS is colored in deep teal. Mg^2+^ ion is shown as a gray sphere, IPP molecules occupying S1 and S2 sites are shown in red and green, respectively. (B) Co-immunoprecipitation of NgBR/DHDDS mutations introduced at the complex interface. HEK293T cells were co-transfected with NgBR-HA and Flag-DHDDS cDNAs; cells were lysed 48 hours post-transfection and immunoprecipitation performed using anti-flag magnetic beads. The lysate was analyzed by western blotting. (C) Limited proteolysis of the core domain was performed by incubating the protein with increasing amounts of thermolysin (0.005, 0.01, 0.02, 0.04, 0.08, 0.16 mg/ml). Untreated protein sample is denoted as (U). (D) Characterization of *cis*-PTase mutants in the C-terminal region of DHDDS using yeast complementation assay. The *nus1*Δ *rer2*Δ *srt1*Δ deletion strain expressing *G.lamblia cis*-PTase from *URA3* plasmid was co-transformed with *MET15* bearing wildtype (WT) NgBR and the *LEU2* plasmid bearing either WT or mutant variants of DHDDS at the C-terminus. Three variants were analyzed including a triple mutation (3A) corresponding to R306A, F313, L317 and two DHDDS truncation Δ256 and Δ289. The cells were streaked onto complete plates (YPD) or synthetic complete medium containing 1% FOA. The Ura3 protein, which is expressed from the *URA3* marker converts FOA to toxic 5-fluorouracil, forcing the cells to loose *G.lamblia cis*-PTase plasmid. Cell growth was monitored over time to assess phenotypic differences.

The crystal structure is consistent with earlier predictions that the central region of the NgBR/DHDDS interface is similar to that observed in prokaryotic UPPS and yeast Nus1 homodimers (supplementary Fig. 4) ^32,47^. Indeed, mutations of several amino acids in the interface weaken the binding between NgBR and DHDDS when the constructs were transfected into HEK293T cells and proteins immunoprecipitated (Fig. 2B). In contrast to a recent modeling study ^23^, the crystal structure reveals that DHDDS’s C-terminus wraps around the protein complex and makes additional contact with NgBR. This structural feature does not appear to be an artifact of crystallization since partial proteolysis of purified NgBR/DHDDS showed that, only upon removal of NgBR, did DHDDS’s C-terminus become susceptible to proteolysis (Fig. 2C), suggesting that access to DHDDS’s C-terminal region is hindered by NgBR. To further study the contribution of DHDDS’s C-terminus to heterodimer formation and enzyme function, two DHDDS truncation mutants (Δ256, Δ289), where part of α7 and the entire α8 helix was deleted, and a triple mutant (R306A/F313A/L317A), which disrupts the packing of the two helices, were generated (supplementary Fig. 2B). Both deletion mutants failed to support growth of *rer2*Δ/*srt1*Δ/*nus1*Δ yeast cells (lacking endogenous genes critical for *cis*-PTase, ^25^, whereas the triple mutant delayed growth (Fig. 2D).

#### The active site of the NgBR/DHDDS complex

Both NgBR and DHDDS subunits are required for a functional *cis*-PTase ^24,25^. Although NgBR, and its yeast homolog Nus1, have a similar fold as DHDDS and members of the homodimeric *cis*-PTase family, they lack multiple key catalytic residues and have a distorted “active site” unable to accommodate substrate. As we recently demonstrated in a functional study with purified human complex, the NgBR subunit contributes to *cis*-PTase activity through its C-terminal -RXG- motif, which is conserved amongst hetero- and homodimeric enzymes ^33^ and this critical -RXG- motif is disordered in the reported homodimeric Nus1 structure ^32^. Here, based on clear electron densities, we modeled the entire C-terminal region of NgBR, as well as two IPP molecules, one (IPP1) occupying the allylic substrate binding site S1, and the other (IPP2) occupying the homoallylic site S2, and a bridging Mg^2+^ ion (Fig. 3A; supplementary Fig. 1B). The C-terminus of NgBR has an extended conformation and traverses the dimeric interface to complete the active site (Fig. 3B): the mainchain amide groups of Leu-291 and Gly-292 of the -RXG- motif form critical hydrogen bonds with homoallylic substrate IPP2’s β-phosphate, while the side chain of Arg-290 interacts with a water molecule that coordinates the Mg^2+^ ion. Gly-292 of the -RXG- motif enables the peptide to make a tight turn over IPP2 and further stabilizes the turn by forming a hydrogen bond with Arg-85, a conserved DHDDS residue that also participates in the binding of the allylic substrate. The function of the -RXG- motif is thus equivalent to that of the P-loop, which binds the β-phosphate group of the allylic substrate FPP.

**Figure 3.**
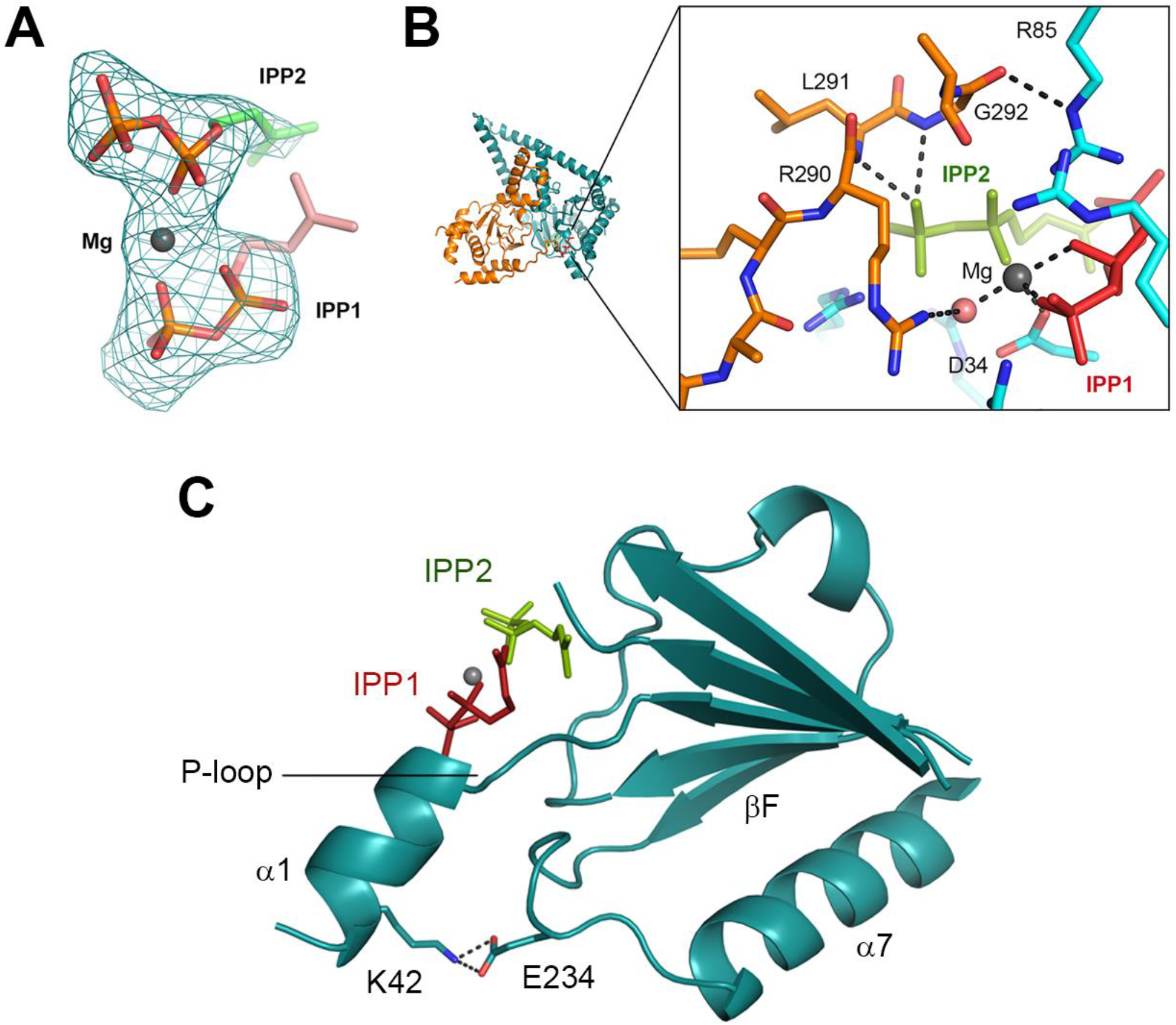
The active site of the NgBR/DHDDS complex. (A) Omit difference map, countered at 3.0 σ level, showing the two IPP molecules and Mg^2+^ ion bound at the active site. IPP1 is assigned to IPP molecule bound at S1 site, and IPP2 to that at S2 site. The oxygen atoms are colored red and phosphorus atoms are colored orange. The carbon atoms of IPP1 are colored salmon and those of IPP2 are in green. (B) Detailed view of the -RXG- motif and the active site. The carbon atoms of NgBR -RXG- motif residues are colored orange and labeled, and those for DHDDS are colored cyan. Nitrogen atoms are colored blue and oxygen atoms are colored red. IPP1 and IPP2 are colored red and green, respectively. Mg^2+^ is shown as a gray sphere, and its co-ordination is indicated by the dashed lines. A coordinating water molecule is shown as red sphere. (C) The K42E retinitis pigmentosa mutation in DHDDS. A cartoon representation indicating the locations of Lys-42 and Glu-234 relative to the P-loop and bound substrates. DHDDS is colored in deep teal, IPP1 in red, IPP2 in green and Mg^2+^ is shown as a gray sphere.

The binding interactions within the S1 site are highly conserved within the *cis*-PTase family (supplementary Fig. 5). Despite weak electron density, we were confident in modeling IPP1 in such a way that its isopentenyl group is pointed toward the hydrophobic tunnel where the elongating product is bound (Fig. 3A). Unlike the S1 site, in previous studies, substrate binding to the S2 site was less consistent among various crystal structures (supplementary Fig. 6) and clear from our study. Some differences were caused by mutations introduced into the active site, but differences may also result from crystallization conditions that destabilized the -RXG- motif since the disordered motif is almost always associated with poorer occupancy of the S2 site, or with misplaced pyrophosphate groups. Amongst known *cis*-PTase structures, the binding mode of IPP2 is most similar to that observed in the complex between MlDPPS and substrate analogs (supplementary Fig. 6). In both structures, the pyrophosphate is tightly bound by three conserved arginine residues (DHDDS Arg-85, Arg-205, Arg-211), a conserved serine (Ser-213), and the -RXG- motif from the dimerization partner. The magnesium ion is coordinated by phosphate groups from both allylic and homoallylic substrates, a conserved aspartate from the “P-loop” (Asp-34), and two water molecules. A conserved asparagine (Asn-82) is identically positioned near IPP’s C2 group for proton extraction during catalysis. These structural similarities reinforce the notion that hetero- and homo- dimeric *cis*-PTases share the same catalytic mechanism.

The majority of CDG-causing missense mutations in DHDDS (R37H, R38H, R211Q) and NgBR (R290H) affects active site residues directly involved in substrate binding and catalysis (supplementary Fig. 7A). Although uncharacterized biochemically, the DHDDS^T206A^ mutation could also perturb the active site because the threonine hydroxyl is simultaneously hydrogen-bonded to the backbone amide and carbonyl of the metal-binding Asp-34 (supplementary Fig. 7B,^49^). The only exception to this pattern is DHDDS^K42E^, which affects ~17% of Ashkenazi Jewish patients diagnosed with retinitis pigmentosa ^42,50^. It was previously unclear how this mutation could affect enzyme activity because the corresponding residue in *E. coli* UPPS (Lys-34) is exposed and does not interact with any other residue that could link the mutation to the active site. Here, we show that instead of pointing toward solvent, Lys-42 forms a salt bridge with the highly conserved Glu-234 (Fig. 3C); and this interaction is equivalent to that between Trp-31 and Asp-223 in *E. coli* UPPS. Therefore, this charge reversal mutation could be disruptive to protein structure, affecting the short helix (α1) that contains Lys-42. Given the role of α1 and the preceding P-loop in FPP binding, this interpretation is consistent with the observation that K42E mutation increases the K_M_ for FPP, decreases k_cat_, but has no effect on IPP binding ^33^. K42E also causes product chain shortening ^25,49^, which will be discussed below.

#### NgBR mutations and Parkinson’s disease

Recently, a number of mutations in the *NUS1* gene were discovered by exome sequencing in patients with Parkinson’s disease (PD) ^45^. Amongst the 15 missense mutations, 11 occur in the region that we now have structural information for (Fig. 4A). In contrast to CDG-causing mutations, which cluster around the active site, most PD mutations are scattered throughout the three-dimensional structure and do not appear to have any direct effect on protein folding, heterodimerization with DHDDS, or substrate binding. Therefore, for these mutations, it remains uncertain whether they altered *cis*-PTase activity *in vivo*, or impacted another biochemical mechanism involving NgBR that is unrelated to *cis*-PTase or dolichol function. It is not yet known whether any of the PD patients had symptoms overlapping with the CDG spectrum earlier in life ^45^, or if CDG patients with mild DHDDS mutations, e.g., K42E, could have a higher risk for developing Parkinson’s disease.

**Figure 4.**
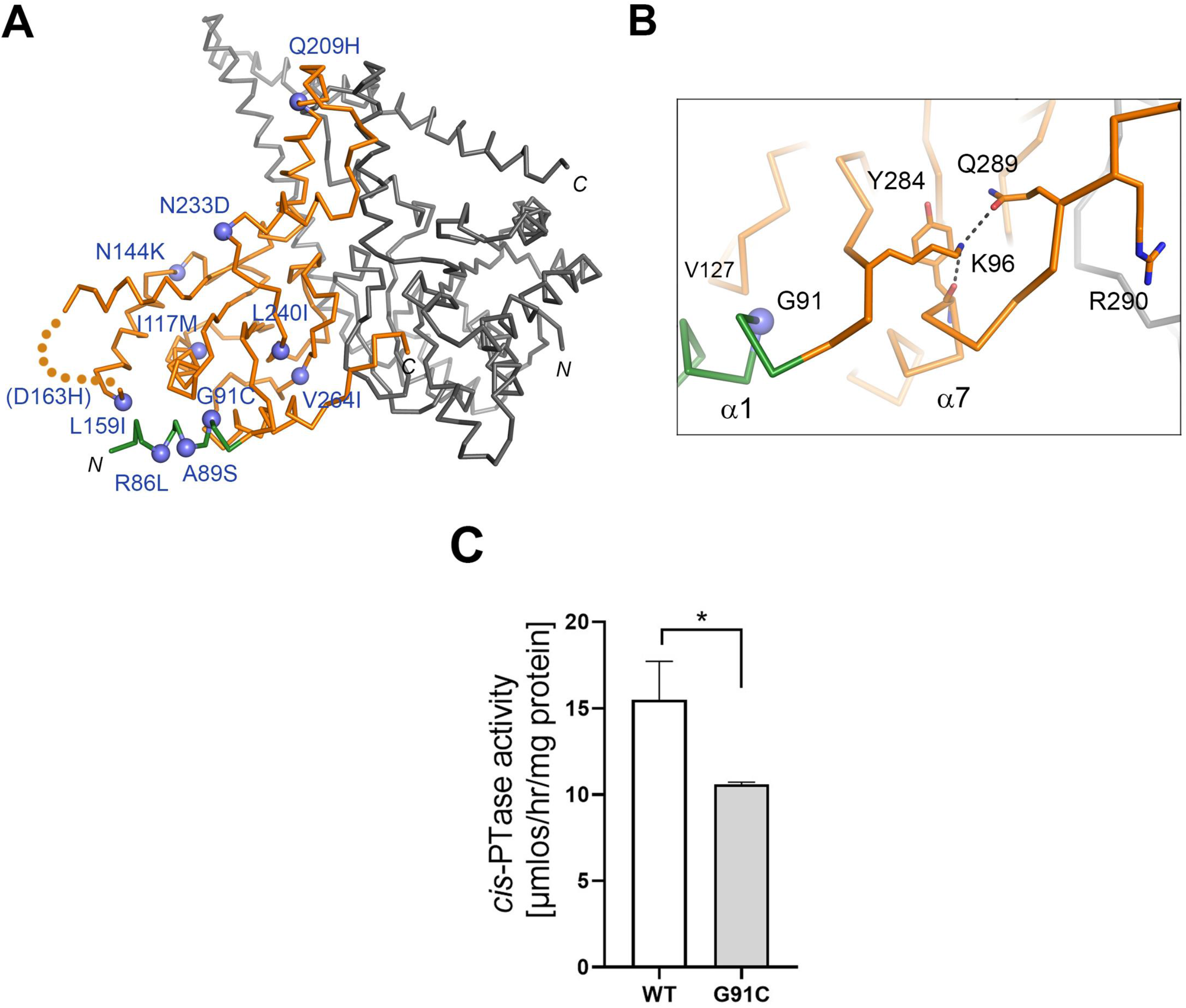
Missense NgBR mutations associated with Parkinson’s disease. (A) NgBR mutations related to Parkinson’s disease are shown as purple spheres and labeled. DHDDS is colored in gray and NgBR is colored in orange except for the N-terminal helix (α1) which is shown in green. (B) Detailed view showing the location of G91C disease mutant within NgBR colored orange except for α1 helix, shown in green. Gly-91 is involved in hydrophobic packing against Val-127. Lys-96 forms hydrogen bonds with Tyr-248 and Gln-289, which stabilize the C-terminal -RXG- motif. (C) *cis*-PTase activity was measured using purified wildtype and NgBR disease mutant, G91C. The mutant exhibits ~ 40% reduction in *cis*-PTase activity compared to wildtype enzyme. The values are the mean ±S.D. of three independent measurements.

One PD mutation (NgBR^G91C^), however, could indirectly affect the enzyme’s active site through the -RXG- motif. As alluded to earlier, the NgBR structure has an N-terminal helix (α1) that is absent in either yeast Nus1 or bacterial UPPS structures (highlighted in green in Fig. 4A). A lysine residue (Lys-96) at the end of this helix plays an important structural role in stabilizing the NgBR’s C-terminal segment by forming two critical hydrogen bonds with the backbone carbonyl of Tyr-284 and the side chain of Gln-289 (Fig. 4B). Gly-91 mediates the contact between α1 and the rest of the protein and introducing a cysteine at this position would be incompatible with the packing of the helix and could perturb, through Lys-96, the C-terminal segment that harbors the -RXG- motif. To investigate this possibility, we generated and biochemically characterized the NgBR^G91C^ mutant. The purified, G91C complex had a ~40% reduction in the enzyme’s specific activity *in vitro* (Fig. 4C). In comparison, the CDG mutation DHDDS^K42E^ causes an 80% reduction in enzyme activity compared to wildtype activity. This finding is significant and raises the possibility that subtle changes of *cis*-PTase activity could contribute to the pathogenesis of PD. These data are consistent with the observation that a splice variant of NgBR reducing its mRNA levels by 50% also increases PD risk ^45^. Dolichol is the most abundant lipid associated with neuromelanin, a dark pigment enriched in catecholaminergic neurons that are selectively vulnerable in Parkinson’s disease ^51,52^. The genetic finding of mutations affecting a critical component of the enzyme complex responsible for dolichol synthesis, and our biochemical characterization of one such mutation, suggest a possible link between dolichol metabolism and age-related neurodegeneration.

#### Regulation of enzyme activity by membrane binding

Heterodimeric *cis*-PTases are invariably found to be peripherally associated with large hydrophobic structures like membrane bilayers, lipid droplets and rubber particles ^12,24,53^. Two hydrophobic segments within the N-terminal region of NgBR is responsible for stable anchoring of the NgBR/DHDDS complex to the ER membrane. Our crystallographic analysis of the core catalytic domain reveals a unique N-terminal structure within DHDDS that is absent in bacterial UPPS. The N-terminal segment consists of a random coil (residues 1-10) and a short α-helix (α0; residues 11-21). The coil starts from the gap between α2 and α3. A bulky side chain (Trp-3) is inserted between the two helices, constricting the hydrophobic tunnel that is predicted to house the polyisoprenyl chain (Fig. 5A, left pane; Fig. 5B). A conformational change displacing the coil from the tunnel is thus required to enable product elongation (Fig. 5A, right panel). The short helix (α0) lies alongside α7 and forms the outermost tip of the entire protein complex. Intriguingly, the side of α0 that is exposed to the solvent contains three hydrophobic residues (Trp-12, Phe-15, Ile-19). Sequence analysis indicates that this continuous hydrophobic patch is conserved in all DHDDSs, raising the possibility that α0 may have evolved specifically for membrane binding (Fig. 5C). We generated a triple DHDDS mutant (W12A/F15A/I19A) to examine the role of the hydrophobic patch in enzyme function. The mutant has similar activity as the wildtype enzyme in the absence of phospholipid. However, W12A/F15A/I19A is no longer potently activated by the addition of phosphatidylinositol (Fig. 5D) and the dominant polyprenol generated by the W12A/F15A/I19A mutant is also three isoprene units longer (Fig 5E). We hypothesize that membrane binding would trigger a conformational change that results in the unblocking of the hydrophobic tunnel during chain elongation and destabilizing the enzyme:product complex to facilitate product release (see below). Therefore, DHDDS’s α0 may function as a membrane sensor, contributing to enzyme activation by phospholipid and to the release of polyisoprenyl product of the appropriate chain length.

**Figure 5.**
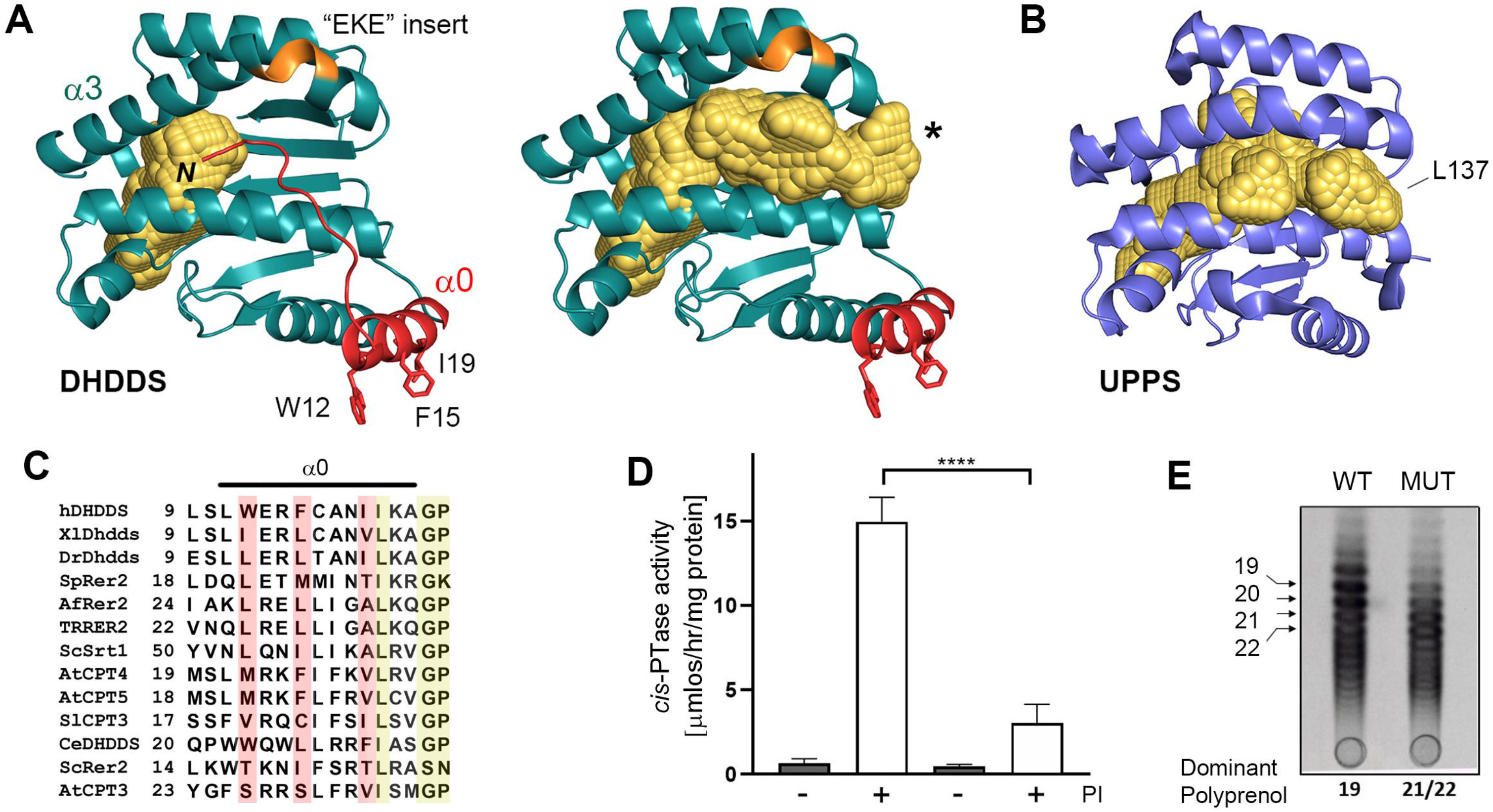
DHDDS’s helix α0 functions as a membrane sensor. (A) A cartoon representation of DHDDS subunit colored in deep teal. The hydrophobic cavities of DHDDS before (left) and after (right) N-terminal loop deletion (residues 1-10) are shown in yellow. The N-terminal loop and α0 helix are shown in red; the side chains of the three exposed hydrophobic residues are shown and labeled. The hydrophobic cavity was generated using the 3V web server ^54^. (B) A cartoon representation of *E.coli* UPPs (PDB entry 1X06) monomer colored in purple. The hydrophobic cavity is shown in yellow. Leu-137 involved in chain length control in UPPS is located at the end of the cavity. (C) Sequence alignment showing the conservation of three hydrophobic amino acids at the N-terminal helix α0 among DHDDS orthologs (highlighted in red). Conserved residues are highlighted in yellow. Proteins represented in this alignment are orthologs of human DHDDS *cis*-PTase subunit as follows: hDHDDS (human, UniProtKB Q86SQ9-1), XlDhdds (*Xenopus laevis;* UniProtKB Q7ZYJ5), DrDhdds (*Danio rerio*, UniProtKB Q6NXA2), CeDHDDS (*Caenorhabditis elegans,* UniProtKB Q5FC21), ScRer2 (*Saccharomyces cerevisiae*, UniProtKB P35196), ScSrt1 (*Saccharomyces cerevisiae*, UniProtKB Q03175), SpRer2 (*Schizosaccharomyces pombe*, UniProtKB O14171), TrRER2 (*Trichoderma reesei,* UniProtKB - G0ZKV6), AfRer2 (*Aspergillus fumigatus* UniProtKB - Q4WQ28), SlCPT3 (*Solanum lycopersicum*, UniProtKB K7WCI9), AtCPT3 (*Arabidopsis thaliana*, UniProtKB Q8S2T1), AtCPT4 (*Arabidopsis thaliana*, UniProtKB Q8LAR7), AtCPT5 (*Arabidopsis thaliana*, UniProtKB Q8LED0) (D) Phospholipid stimulation of wildtype and W12A/F15A/I19A triple mutant is shown. Stimulation was compared by measuring *cis*-PTase activity of purified wildtype and DHDDS triple mutant in the presence and absence of 1% phosphatidylinositol (PI). The values are the mean ±S.D. of five to eight independent measurements. (E) Reverse phase TLC separation of dephosphorylated products from *cis*-PTase activity of WT and DHDDS triple mutation (W12A/F15A/I19A) denoted as MUT. Numbers correspond to the dominant polyprenols in each sample are shown at the bottom of the plate. The position of the polyprenol standards is shown on the left.

#### The mechanism determining product chain length

Eukaryotic heterodimeric *cis*-PTases differ categorically from bacterial homodimeric enzymes in that they not only generate products with longer chain lengths but also tend to generate a range of products. The human NgBR/DHDDS complex preferentially synthesizes C_95_, but as Fig. 5E illustrates, C_95_ is one of many polyprenols that differ by single isoprene unit and are present in varying amounts. Here we examine the mechanistic basis for this biochemical phenomenon in light of the new crystal structure.

The size of the hydrophobic tunnel cannot explain the longer product (C_95_) generated by the NgBR/DHDDS complex. After manually removing the N-terminal coil that blocks the hydrophobic tunnel (Fig. 5A), we used 3V web server to calculate the volume of DHDDS’s tunnel to be around 1,300 Å^3^ ^54^, which is comparable in size to that (1,200 Å^3^) of *E. coli* UPPS (Fig. 5B; UPPS generates C_55_ product). Therefore, as polyprenol intermediates are formed, either the protein tunnel has to expand significantly, or the tail of the polyprenol chain needs to be pushed out of the tunnel. A model of protein conformational change that involves a large movement of helix α3 has been proposed ^55^. This type of movement, however, is unlikely to occur during product elongation because α3’s N-terminal half is involved in binding the allylic substrate’s pyrophosphate group, and thus essential for subsequent rounds of reaction. We favor the second possibility where a large portion of the polyprenol chain exits the active site tunnel and becomes exposed on the protein surface. Mutagenesis studies performed on *E. coli* UPPS suggested that Leu-137, equivalent to DHDDS’s Cys-148, located near the end of the tunnel could function as the barrier to such an exit since the UPPS^L137A^ mutant synthesizes a range of long-chain products (up to C_75_) in the absence of detergent (Fig. 5B) ^19^.

Sequence alignment revealed that all eukaryotic heterodimeric *cis*-PTases carry a short insertion (human DHDDS Glu-107, Lys-108, Glu-109), in the middle of the α3 helix (Fig. 6A). The extra sequence creates a greater bulge in α3 but does not significantly increase the volume of the hydrophobic tunnel that lies beneath the helix (Fig. 5A; left panel and Fig. 6C). To examine the possibility that the insertion may impact product length, we generated an ΔEKE deletion mutant of DHDDS and transformed it and wildtype DHDDS into the *S. cerivisiae* triple deletion strain (*rer2Δ/srt1Δ/nus1Δ*) with wildtype NgBR. As expected, when expressed in yeast, the wildtype enzyme generated Dol-20 (C_100_) as the dominant product. Although the ΔEKE mutant generated shorter products (Fig. 6B), it retained the two key characteristics of long chain *cis*-PTase (i.e. chain length distribution of products and each product > C_55_). Therefore, the wider opening between α2 and α3 created by the sequence insert, *per se*, is not a required structural feature of the long-chain enzyme.

**Figure 6.**
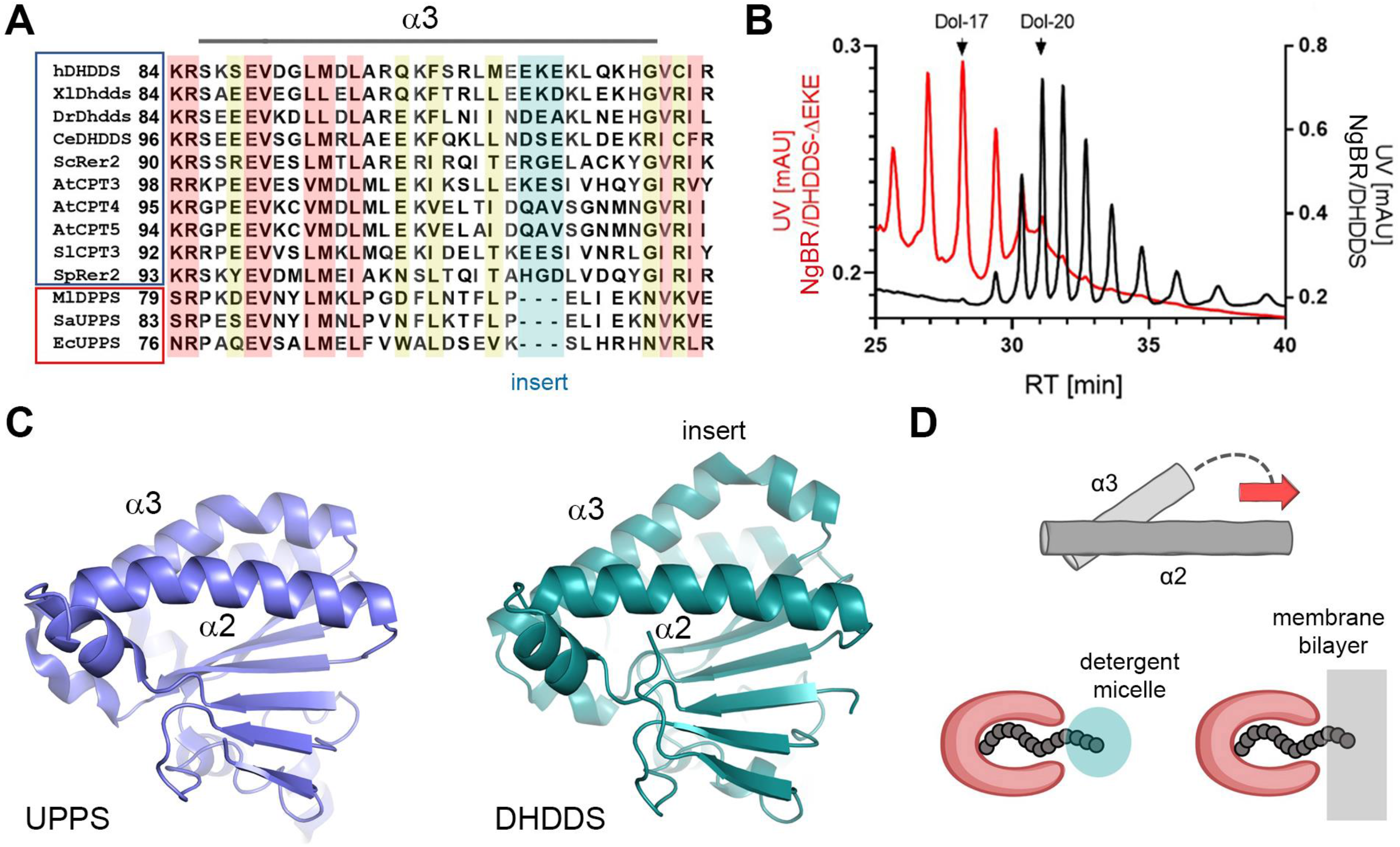
Mechanism of chain elongation by heteromeric *cis*-PTases. (A) Multiple amino acid sequence alignment comparing α3 helix between DHDDS orthologs and homodimeric *cis*-PTases. Highly conserved residues are highlighted in red and less conserved ones are shown in yellow. Region corresponding to DHDDS EKE insert is highlighted in blue; the region is missing in homodimeric enzymes and a gap is present instead. Proteins represented in this alignment are: single subunit *cis*-PTs: EcUPPS (*Escherichia coli,* UniProtKB P60472), MlDPPS (*Micrococcus luteus*, UniProtKB O82827), SaUPPS (*Sulfolobus acidocaldarius,* UniProtKB Q9HH76). Orthologues of human DHDDS *cis*-PTase subunit: hDHDDS (human, UniProtKB Q86SQ9-1), XlDhdds (*Xenopus laevis,* UniProtKB Q7ZYJ5), DrDhdds (*Danio rerio,* UniProtKB Q6NXA2), CeDHDDS (Caenorhabditis elegans, UniProtKB Q5FC21), ScRer2 (*Saccharomyces cerevisiae*, UniProtKB P35196), SpRer2 (*Schizosaccharomyces pombe*, UniProtKB O14171), SlCPT3 (*Solanum lycopersicum*, UniProtKB K7WCI9), AtCPT3 (*Arabidopsis thaliana*, UniProtKB Q8S2T1), AtCPT4 (*Arabidopsis thaliana*, UniProtKB Q8LAR7), AtCPT5 (*Arabidopsis thaliana*, UniProtKB Q8LED0) (B) HPLC analysis of chain length of dolichol generated by WT and ΔEKE DHDDS mutant in yeast cells. The dolichol peaks were identified and labeled on top of the chromatogram. The WT cells yielded the main compound Dol-20 compared to Dol-17 in ΔEKE mutant. The chain length and identity of lipids were confirmed by comparison with external standards of a polyprenol (Pren-10 – Pren-24) and dolichol (Dol-17 – Dol-23) mixtures. (C) Structural comparison between *E. coli* UPPS (PDB ID 1X06) monomer and human DHDDS subunit. The EKE insert within α3 helix of DHDDS creates a bigger bulge that may contribute to the stabilization of the enzyme:product complex during chain elongation. (D) Schematic diagram illustrating the proposed chain elongation mechanism for *cis*-PTases. Red arrow indicates the direction of product elongation. Exposed hydrophobic isoprene units may increasingly destabilize the enzyme:product complex by interacting with detergent micells (blue) or lipid bilayers (grey).

If the tail of the elongating polyprenol product does exit the enzyme’s hydrophobic tunnel, this is expected to generate several consequences (Fig. 6D). (i) Since there is no longer a physical barrier, the polyprenol chain can grow indefinitely in theory. This distinguishes the long-chain *cis*-PTases from the short-chain and medium-chain *cis*-PTases. In the latter two, the product chain length is strictly determined by the size of the hydrophobic tunnel ^19,21^. (ii) The stability of the enzyme complex with product intermediates likely plays a major role in determining when elongation reaction stops. The exposed portion of the hydrophobic polyprenol could be increasingly disruptive to the enzyme:product complex through various mechanisms, e.g., interaction with detergent micelles, membrane bilayers, or lipid droplets. In the example cited above, *E. coli* UPPS^L137A^ mutant failed to produce long-chain product in the presence of TX100 ^19^. (iii) The insert in α3, we thus argue, plays a role in stabilizing the enzyme:product complex. One possibility, as schematically illustrated in Fig. 6D, is that the insert may confer greater conformational flexibility to the α3 helix and its surrounding regions. (iv) The subtle effect of the environment on the enzyme:product complex determines that the final product will invariably have a distribution of chain lengths, another defining feature of the long-chain *cis*-PTases. The most extreme example of a long chain *cis*-PTase is rubber synthase, together with its protein cofactors, must have extraordinary stability in its product-bound form ^12^.

In summary, we provide the first atomic insights into how eukaryotes synthesize dolichol, the obligate carrier lipid for N-glycosylation, O- and C-mannosylation reactions and GPI anchor biosynthesis. The new features elucidated by this structure rationalize the stability of the heterodimeric complex, disease causing mutations and lipid regulation of *cis*PTase activity. The mechanism suggested by the NgBR/DHDDS crystal structure provides a conceptual framework for understanding the unique enzymatic properties of the long-chain *cis*-PTases, including rubber synthase. Obligate heterodimerization with a membrane-binding partner probably ensures that the long-chain product is only robustly synthesized in close vicinity to the membrane as elsewhere the exposed polyprenol could be harmful to the cell.

## Acknowledgements

This work was supported by Grants R35 HL139945 (to WCS), an American Heart Association Scientist Development Grant (to EJP) and a National Science Centre of Poland Grant UMO-2016/21/B/NZ1/02793 (to LS). Additionally, the work was supported by the Northeastern Collaborative Access Team beamlines, which are funded by the National Institute of General Medical Sciences from the National Institutes of Health (P30 GM124165). The Pilatus 6M detector on 24-ID-C beam line is funded by a NIH-ORIP HEI grant (S10 RR029205). This research used resources of the Advanced Photon Source, a U.S. Department of Energy (DOE) Office of Science User Facility operated for the DOE Office of Science by Argonne National Laboratory under Contract No. DE-AC02-06CH11357.

## Author contributions

BE, KG, EJP, RZ, YA and WCS designed experiments; BE, EJP, KG, BS performed experiments; BE and RZ conducted protein purification and crystallization trials; BE and YH analyzed crystallographic data; LS performed HPLC analysis of dolichol species; BE, KG, YH and WCS wrote the manuscript. All authors contributed to the review of the manuscript. YH and WCS are co-corresponding authors. The project emanated from the Sessa lab and where the proteins were purified, and enzymology studied. The Ha lab was essential for crystallization expertise and structural insights resulting in a logical arrangement of both being co-corresponding authors.

## EXPERIMENTAL PRECEDURES

### Materials

Unless otherwise stated, all reagents were of analytical grade and purchased from Sigma-Aldrich, Thermo Fisher Scientific, and Zymo Research (Irvine, CA). Restriction enzymes were from New England Biolabs (Ipswich, MA). [1-^14^C] IPP (50 mCi/mmol) was purchased from American Radiolabeled Chemicals (St. Luis, MO). Reverse phase thin layer chromatography (RP18-HTLC) plates were from MilliporeSigma (cat# 1.51161.0001). Primary antibodies used in this study include Anti-HA High Affinity antibody (Roche, 11867423001) and Monoclonal anti-Flag M2 antibody (Sigma, F3165). HiFi DNA Assembly method (NEBuilder^®^, NEB) was used to construct expression vectors and perform site-directed mutagenesis. List of plasmids is in supplementary Table 2–4. Primers used in cloning are listed in supplementary Table 5 and primers used for mutagenesis are listed in supplementary Table 6.

### Cloning and Purification of NgBR/DHDDS Complex

To express His-SUMO-NgBR^79-293^ and untagged, full length DHDDS (1-333) in bacteria, His-SUMO and NgBR overlapping PCR fragments were first assembled into pRSF-DUET1 vector cut with NdeI/XhoI restriction enzymes. DHDDS PCR fragment was then assembled into pRSF-DUET1-HIS-SUMO-NgBR cut with NcoI/NotI. The Recombinant NgBR/DHDDS complex was expressed in *Escherichia coli* Rosetta (DE3) cells (Novagen) and induced with 0.7 mM IPTG (OD_600_ 0.6) overnight at 18 °C. Cells were harvested and then resuspended in lysis buffer containing 20 mM Tris-HCl pH 8.0, 500 mM NaCl, 20 mM imidazole, 10% glycerol, 0.5% triton X-100 and 2 mM 2-Mercaptoethanol, cOmplete protease inhibitors (Roche), lysozyme (100 μg/ml) and DNase I (10 μg/ml). Three cycles of freeze/thaw were conducted using ethanol/dry ice bath and cells were sonicated in 50 ml falcon tube for 2 minutes total. The samples were clarified by centrifugation at 20,000 rpm for 1 hour at 4 °C. Supernatant was then applied to a 1 ml HisTrap (GE Healthcare) nickel affinity column, and the protein was eluted with 6 ml lysis buffer containing 400 mM imidazole. The sample was then applied to size exclusion column (Superdex 200, GE Healthcare) equilibrated with buffer containing (50 mM Tris-HCl pH 8.0, 150 mM NaCl, 1 mM MgCl_2_ and 2 mM TCEP). Fractions containing protein complex were collected and subjected to cleavage with SUMO protease overnight at 4°C to remove the His-SUMO tag. The cleaved protein was then re-applied to a HisTrap column and the flow-through was collected. Protein sample was then passed through another size exclusion column equilibrated with 50 mM Tris-HCl pH 8.0, 150 mM NaCl, 2.5 mM MgCl_2_ and 2 mM TCEP. Fractions were collected and analyzed by 12% SDS-PAGE gel.

### Crystallization, Data Collection and Structural Determination

The purified protein was concentrated to 3.2 mg/ml and incubated with 3.3 mM IPP (sigma) on ice for 2 hours. Crystallization screening was performed using the sitting-drop vapor diffusion method, and an initial hit was obtained from PEG screen (Hampton Research). Crystallization was optimized by grid screening and the best crystals were obtained by mixing 1 μl protein solution with 1 μl reservoir solution consisting of 0.1 M Bicine (pH 8.5), 10% v/v 2-propanol, 22% PEG 1500. Crystals appeared within 2 days and grew to maximum size within one week at room temperature. Crystals were cryoprotected with the reservoir solution supplemented with 20% glycerol and flash frozen in liquid nitrogen. Diffraction Data were collected on beamline 24-ID-E of the Advanced Photon Source at Argonne National Laboratory and processed using *HKL2000* ^56^. The structure of the complex was determined by molecular replacement and refined using CCP4i (supplementary Table 1) ^57^. The *E.coli* UPPs (PDB entry 1X06) and *S.cerevisiae* NUS1 (PDB entry 6JCN) were used as search probes for the DHDDS and NgBR subunits, respectively. Model building were performed using *Coot ^58^*.

### *cis*-PTase activity of NgBR/DHDDS

The steady state activity of purified NgBR/DHDDS complex was assayed as before with minor modifications ^33^. Briefly, a standard incubation mixture contained, in a final volume of 25 μl, 100 μM [1-^14^C] IPP, 20μM FPP, 50 mM Tris-HCl pH 8.0, 1 mM MgCl_2_, 10 mM KF, 20 mM 2-mercaptoethanol, 1 mg/ml BSA ,1 % Phosphatidylinositol and 100 ng of purified enzyme. The mixture was incubated for 1 hour at 37 °C and product was extracted with chloroform:methanol (3:2), followed by washing three times with 1/5 volume of 10 mM EDTA in 0.9% NaCl. The incorporation of [^14^C] IPP into organic fractions containing polyprenol pyrophosphate was measured by scintillation counter.

In order to determine the chain length of the cis-PTase products, polyprenol diphosphates were chemically dephosphorylated by incubation of the lipids at 90° in 1 *N* HCl for 1 hr. Dephosphorylated prenols were extracted three times with two volumes of hexane. The organic fraction was washed with 1/3 volume of water, hexane was evaporated under stream of nitrogen and lipids were loaded onto HPTLC RP-18 precoated plates and run in acetone containing 50 mM H_3_PO_4_. The plates were exposed to film to visualize the products of IPP incorporation. As an internal and external standards Geranylgeraniol (Echelon Biosciences), Undecaprenol and Polyprenol 19 (*Institute of Biochemistry and Biophysics*, PAS the *Collection* of Polyprenols) and Prenols mixture (13-21) (Avanti Polar Lipids) were used. Prenol standards were visualized by exposing the TLC plate to iodine vapor.

### Limited Proteolysis

20 ul of 0.2 mg/ml of purified NgBR/DHDDS enzyme was incubated with 5 ul of Thermolysine (*sigma*) at different concentrations (0.005, 0.01, 0.02, 0.04, 0.08, 0.16 mg/ml). The reaction mixture was incubated at room temperature for 30 minutes and stopped with SDS buffer containing 5.2 mM PMSF and 5.2 mM EDTA. Samples were boiled for 5 minutes and analyzed by12% SDS-PAGE gel.

### Co-Immunoprecipitation

HEK293T cell was transfected using lipofectamine 2000 (Invitrogen) according to the manufacturer’s protocol and harvested 48 hrs after transfection. Cells were collected and lysed in IP buffer (IP buffer: 50mM HEPES, 150mM NaCl, 1mM EDTA, 1% Triton X-100, Complete Protease Inhibitors (Roche)). Lysates were cleared by centrifugation at 12000 rpm for 10 min and 10μl of anti-Flag M2 magnetic beads (Sigma) was used to pulldown the Flag-tagged protein from 0.5-1 mg of cell lysate. After incubation for 2 hours at 4°C, magnetic beads were washed with IP buffer, resuspended in 2X Laemmli sample buffer and boiled for 5 min before western blot analysis.

### Yeast Complementation Assay

For yeast complementation analysis of *cis*-PTase, *S.cerevisiae* strains KG405 (*nus1*Δ *rer2*Δ *srt1*Δ), carrying the *Glcis*-PTase gene on a plasmid with a *URA3* marker was used ^25^. To phenotypically analyze human *cis*-PTase mutants, strain KG405 was transformed with vectors pKG-GW1 carrying DHDDS variants (leucine selection) and pKG-GW2 carrying NgBR variants (methionine selection) in combination or empty vectors as negative control. Transformed yeast cells were grown overnight at 30 °C in synthetic defined medium or lacking uracil, methionine, and leucine were streaked onto synthetic defined medium containing all amino acids, nucleotide supplements, and 1% (w/v) 5-FOA (Zymo Research) and onto YPD plates. The plates were incubated for up to 5 days at 30 °C. Colonies growing on the 5-FOA plates were streaked on synthetic defined medium lacking uracil and incubated at 30 °C for 3 days to verify the loss of the pNEV-Gl*cis*-PTase plasmid. Yeast strain KG405 and its derivative carrying NgBR/DHDDS complex were cultured in 2% (w/v) Bacto peptone and 1% (w/v) yeast extract supplemented with 2% glucose (w/v) (YPD). Synthetic minimal media were made of 0.67% (w/v) yeast nitrogen base and 2% (w/v) supplemented with auxotrophic requirements. For solid medium, agar (BD Biosciences, Sparks, MD) was added at a 2% (w/v) final concentration. Yeast cells were transformed using the Frozen-EZ yeast transformation II kit (Zymo Research).

### HPLC Analysis

To estimate the chain length of dolichols produced by EKE deletion mutant, total lipids from 3 g of yeast cells grown overnight till late logarithmic phase of growth (OD 3-4) were extracted by the modified Folch method. Lipids extracted from yeast cells were hydrolyzed in hydrolytic solution containing toluene/7.5% KOH/95% ethanol (20:17:3, v/v/v) for 1h at 90°C. Nonsaponifiable lipids were then extracted four times with hexane, purified on silica gel 60 columns using isocratic elution with 10% diethyl ether in hexane, evaporated to dryness in a stream of nitrogen and dissolved in isopropanol. Extracts were analyzed by HPLC using a Waters dual-pump HPLC device coupled with a Waters Photodiode Array Detector (spectrum range: 210 - 400 nm) and ZORBAX XDB-C18 (4.6 × 75 mm, 3.5 μm) reversed-phase column (Agilent, USA). Polyisoprenoids were eluted using the solvent mixtures A - methanol/water, 9:1 (v/v) and B - methanol/isopropanol/hexane, 2:1:1 (v/v/v) combined as follows from 0% B to 75% B in 20 min, from 75% B to 90% B in 5 min, from 90% B to 100% B in 2 min, 100% B maintained for 11 min, then from 100% B to 0% B in 1min at a flow rate of 1.5 mL/min. The chain length and identity of lipids were confirmed by comparison with external standards of a polyprenol (Pren-10 – Pren-24) and dolichol (Dol-17 – Dol-23) mixtures.

## Supplementary Figure Legends

**Supplementary Figure 1.**
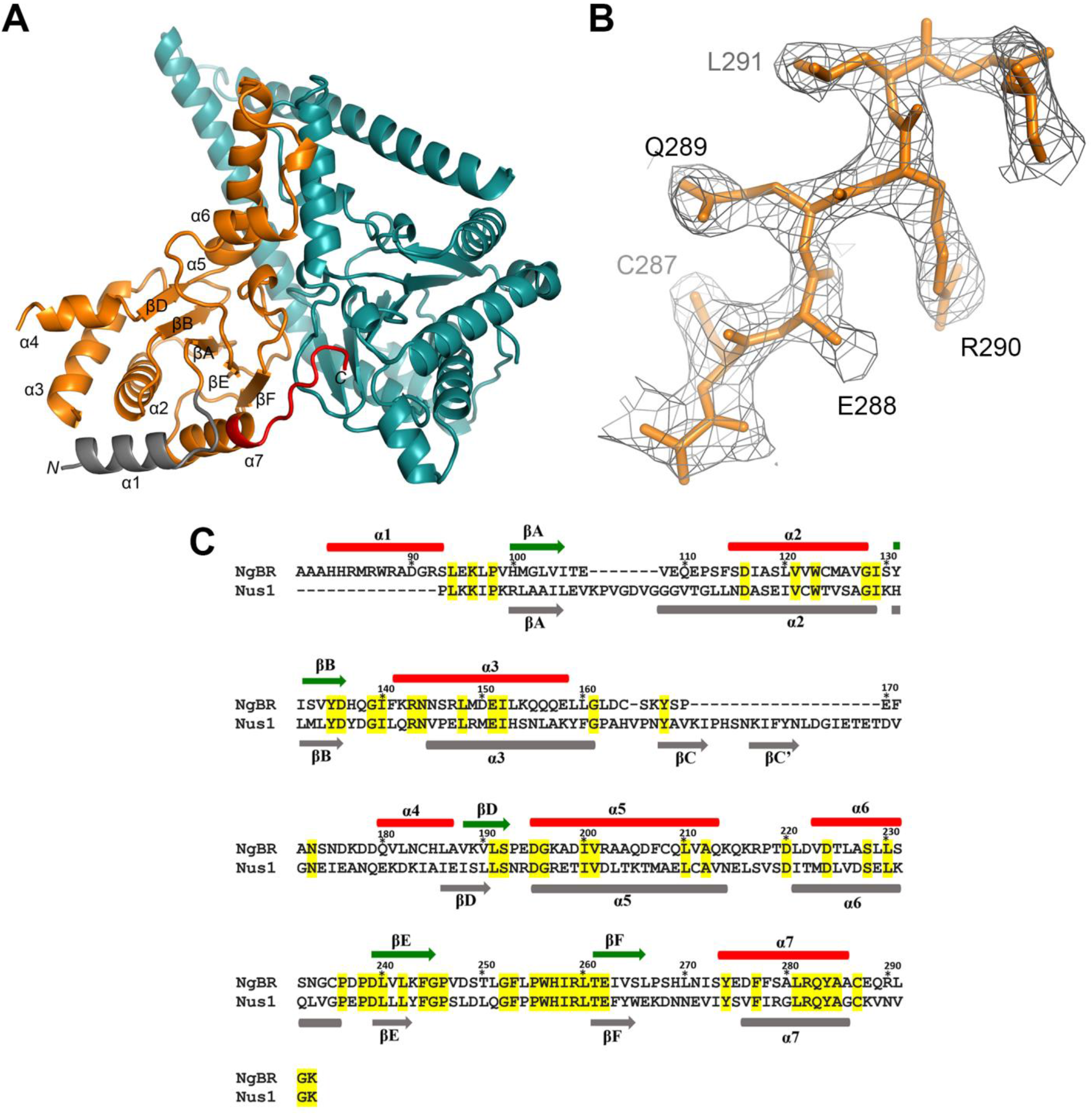
Structural details of the NgBR subunit. (A) Ribbon diagram of NgBR/DHDDS complex; NgBR colored in orange and DHDDS in deep teal. The N-terminal helix of NgBR is colored in gray and the C-terminal loop in red. Secondary structure elements as well as N- and C-termini of NgBR are labeled on the structure. (B) 2Fo-Fc electron density map of the C-terminal -RXG- motif of NgBR contoured at 1.0 σ. The motif residues are colored orange and labeled. (C) Partial amino acid sequence of NgBR subunit (residues 79-293) is aligned with partial sequence of yeast NUS1 subunit (residues 148-375). Numbers and secondary structure elements shown above the sequence correspond to that of NgBR. NUS1 secondary structure elements are shown in gray, below the sequence. Conserved residues are highlighted in yellow.

**Supplementary Figure 2.**
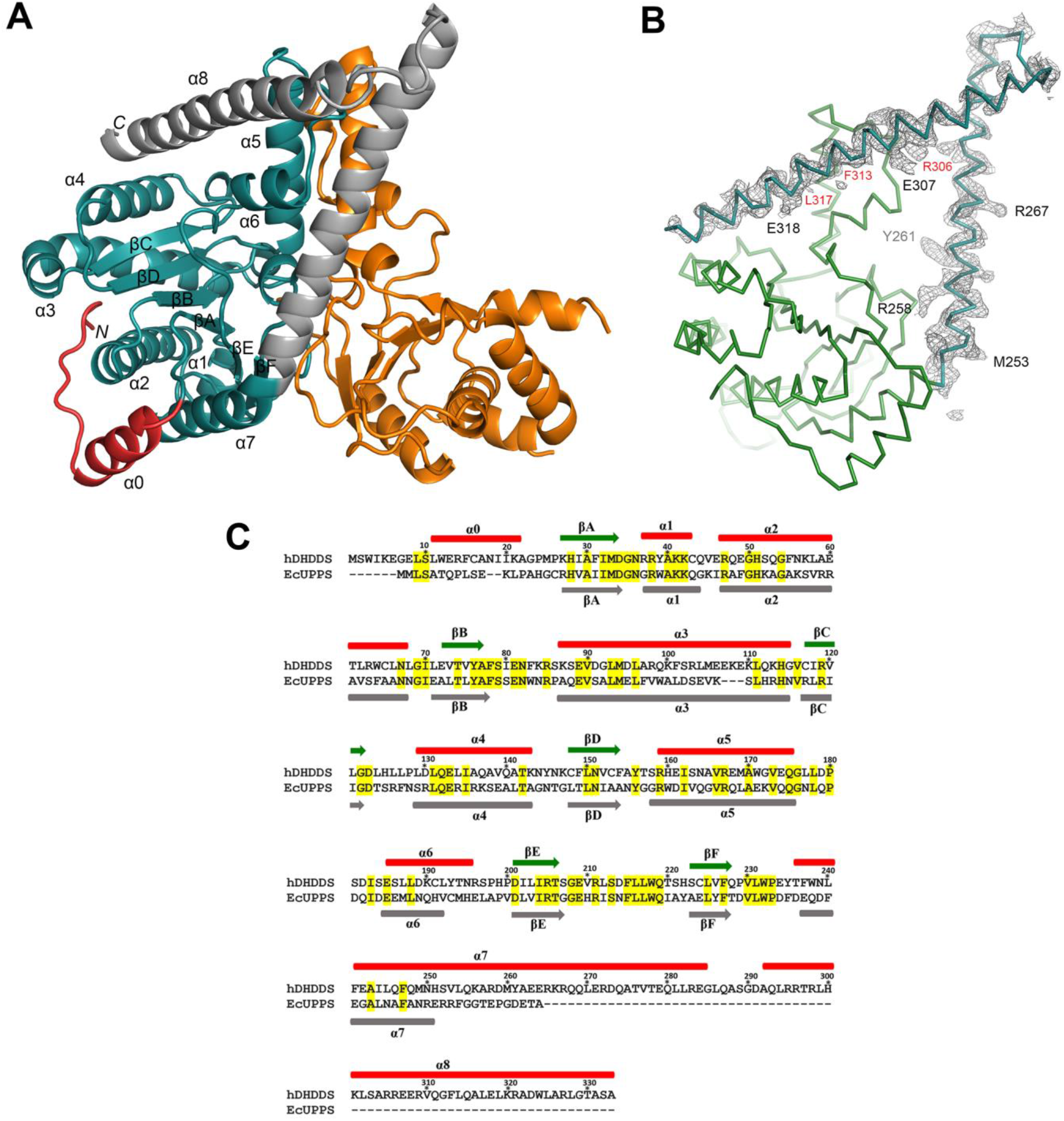
Structural details of the DHDDS subunit. (A) Ribbon diagram of NgBR/DHDDS complex; NgBR colored in orange, DHDDS in deep teal. The N-terminal region of DHDDS is colored red and the C-terminal helices are in gray. Secondary structure elements as well as N- and C-termini of NgBR are labeled on the structure. (B) 2Fo-Fc electron density map of the C-terminal helices of DHDDS contoured at 1.0 σ. Residues mutated in the yeast complementation experiment are labeled in red. (C) The full-length amino acid sequence of DHDDS subunit (residues 1-333) is aligned with full length *E.coli* UPPS (residues 1-253).). Numbers and secondary structure elements shown above the sequence correspond to that of DHDDS. *E.coli* UPPS secondary structure elements are shown in gray, below the sequence. Conserved residues are highlighted in yellow.

**Supplementary Figure 3.**
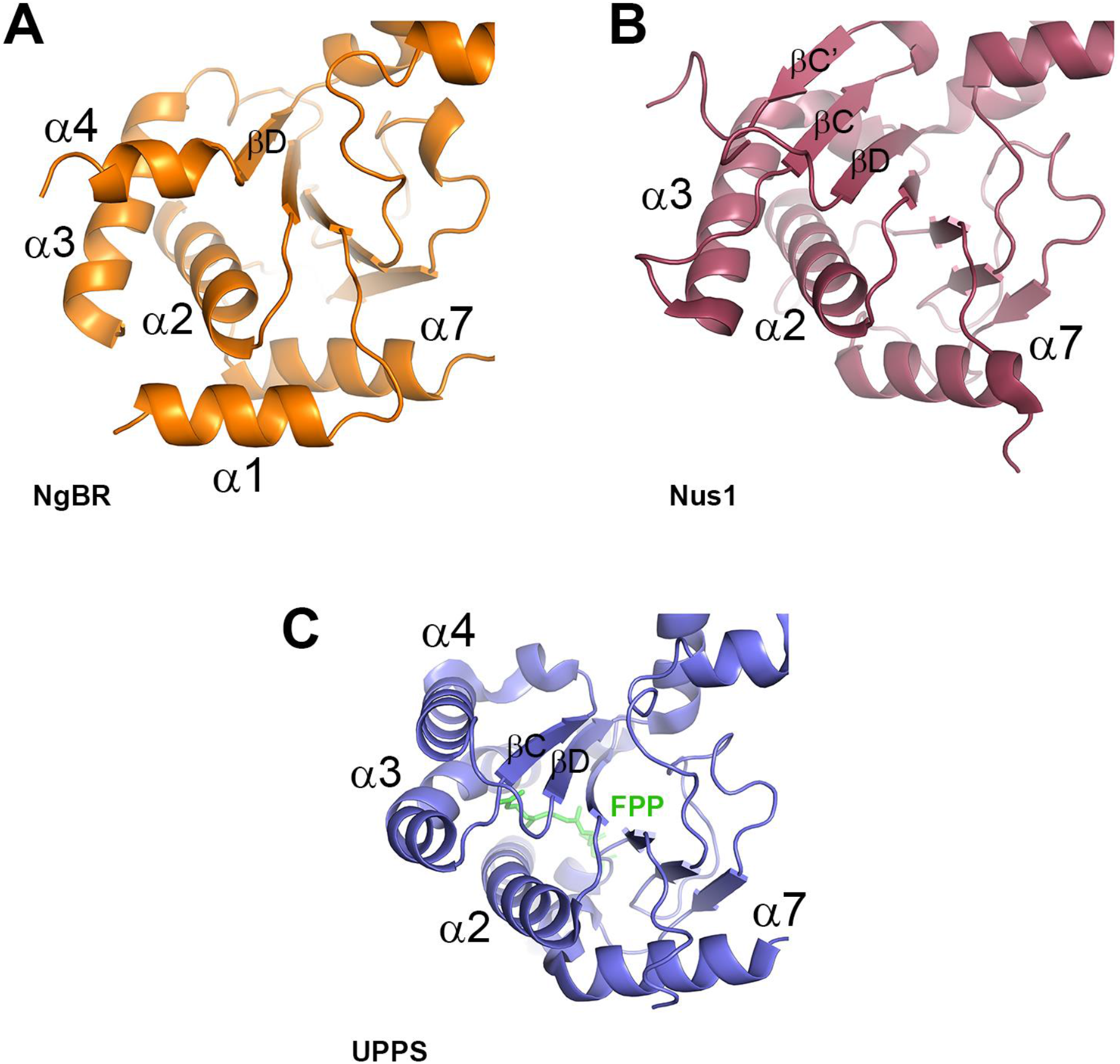
Structural comparison of NgBR, Nus1 and UPPS. A ribbon diagram comparing the structure of (A) NgBR (orange) to that of (B) yeast Nus1 (Ruby, PDB ID 6JCN) and (C) UPPS (Purple, PDB ID 4H8E). NgBR possesses α4 helix, which differs from that of UPPS and is replaced by βC and βC’ in Nus1. An additional α1 helix is present in NgBR N-terminal region and is not observed in Nus1.

**Supplementary Figure 4.**
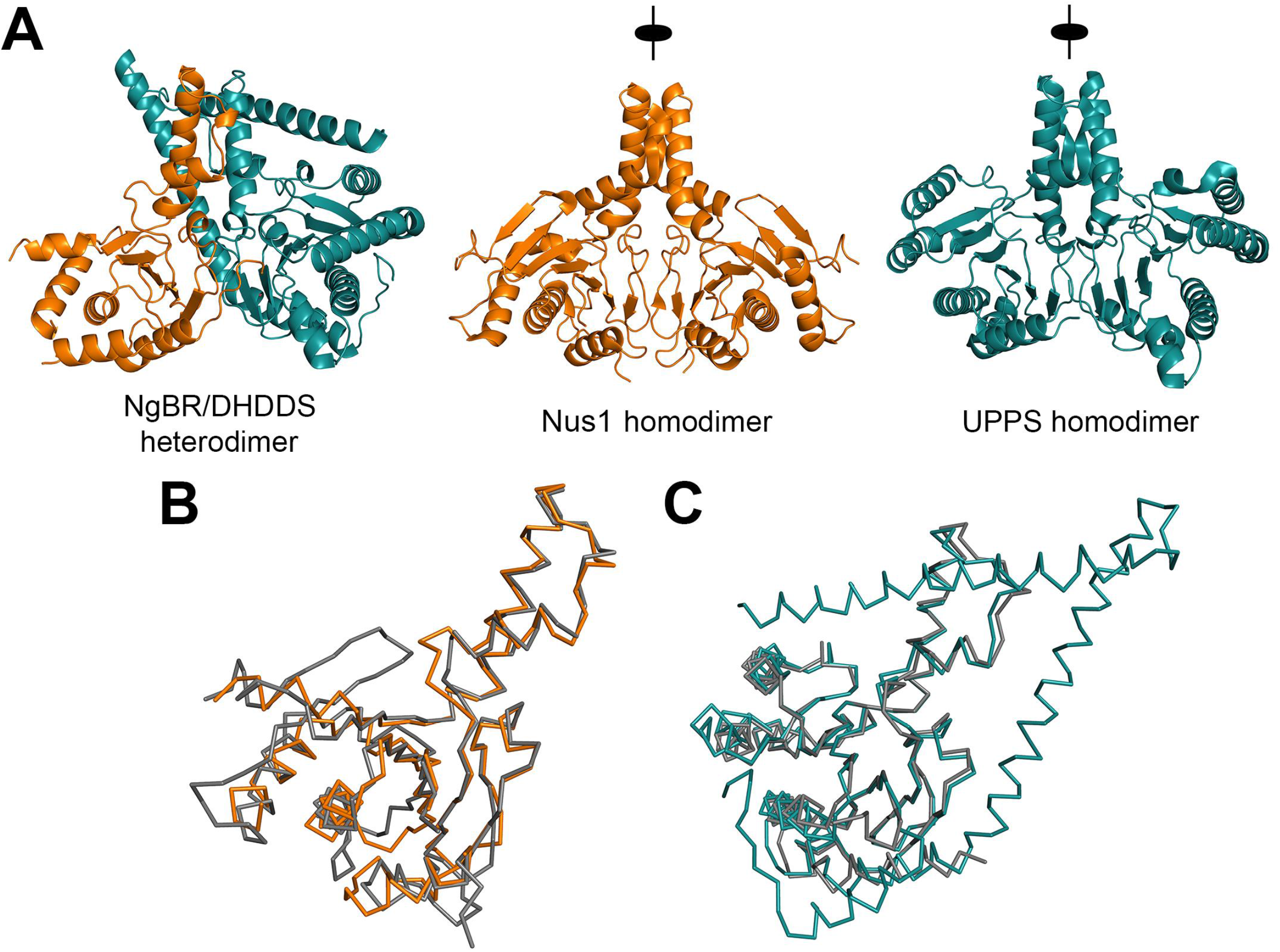
Comparison of different dimeric structures. (A) Ribbon diagram of the heteromeric complex is shown. NgBR is colored in orange, and DHDDS is colored in deep teal. The heteromeric structure is compared with those of yeast NUS1 dimer (PDB 6JCN) colored in orange and *E. coli* UPPS (PDB 1X06) colored in deep teal. (B) NgBR (orange) is superimposed on yeast NUS1 (gray). Wireframe diagram of C_α_ atoms is shown. (C) DHDDS (deep teal) is superimposed on *E. coli* UPPS (gray).

**Supplementary Figure 5.**
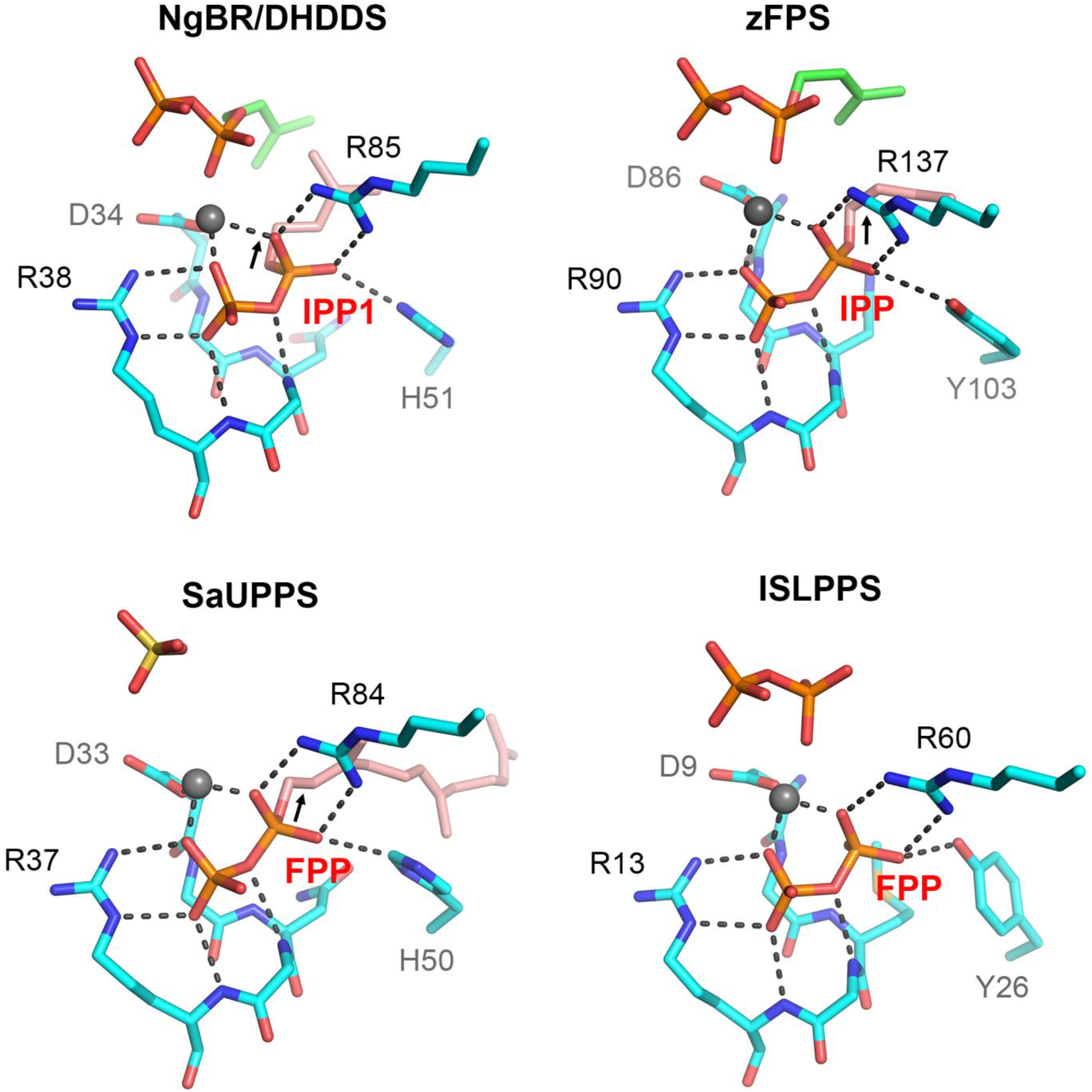
Structural comparison of the S1 site. The S1 site involved in the binding of allylic substrate is compared among different *cis*-PTases including human NgBR/DHDDS complex, *cis*-PTase from tomato, denoted as zFPS (PDB ID 5HXP), *S.aureus* UPPS denoted as SaUPPS (PDB ID 4H8E) and isosesquilavandulyl diphosphate synthase denoted as ISLPPS (PDB ID 5XK6). Mg^2+^ ion is shown as gray sphere, conserved residues involved in binding of allylic pyrophosphate are colored in cyan and labeled, and hydrogen bonds are shown as black-dashed lines. Oxygen atoms are colored red, phosphorus atoms are in orange and nitrogen are in blue. The carbon atoms of the allylic substrate are shown in salmon and those for the homoallylic substrate are shown in green. His-51 in the human NgBR/DHDDS complex is sometimes replaced with tyrosine; in both cases, the side chain is involved in hydrogen bonding with α phosphate of the allylic substrate. His-50 in SaUPPS can also participate in hydrogen bonding if the imidazole ring is rotated 180 degrees.

**Supplementary Figure 6.**
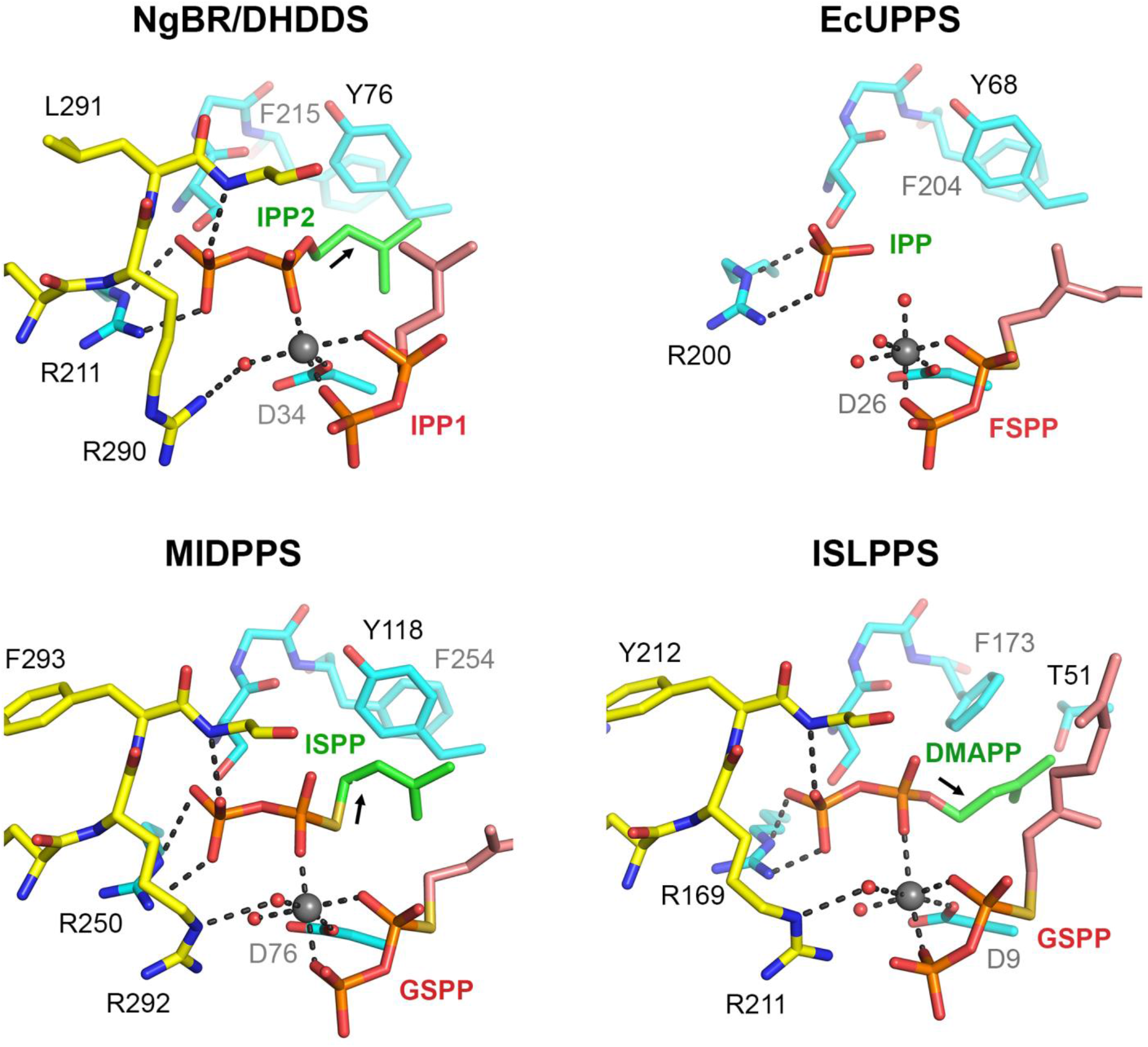
Structural comparison of the S2 site. The S2 site involved in the binding of homoallylic substrate is compared among different *cis*-PTases including human NgBR/DHDDS complex, *cis*-PTase from *E.coli* UPPS, denoted as EcUPPS (PDB ID 1X06), *M.tuberculosis* UPPS Rv2361c denoted as MlDPPS (PDB ID 6IME) and isosesquilavandulyl diphosphate synthase denoted as ISLPPS (PDB ID 5XK9). Mg^2+^ ion is shown as gray sphere, conserved C-terminal -RXG- residues from NgBR and other corresponding monomers are colored in yellow while those for DHDDS are in cyan. Hydrogen bonds are shown as black-dashed lines. Oxygen atoms are colored red, phosphorus atoms are in orange and nitrogen atoms are in blue. The carbon atoms of the allylic substrate are shown in salmon and those for the homoallylic substrate are shown in green. The disordered -RXG- motif in EcUPPS results in an incomplete active site and misplaced IPP phosphate.

**Supplementary Figure 7.**
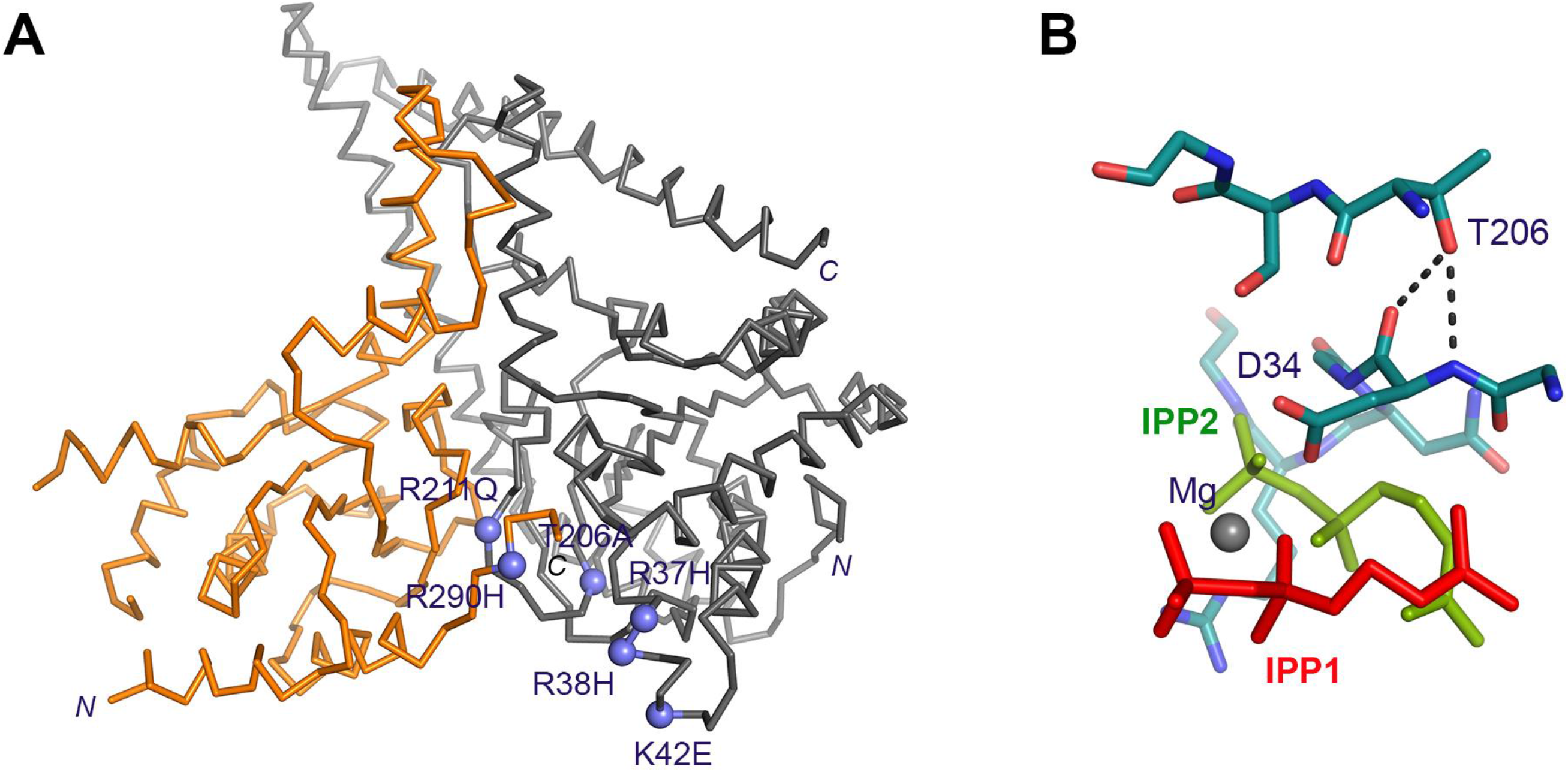
Missense mutations associated with CDG. (A) CDG-causing NgBR and DHDDS mutations are indicated as purple spheres on the structure. NgBR is colored in orange and DHDDS in gray. The CDG mutations cluster around the active site of the complex. In DHDDS, those include: Arg-37, Arg-38, Lys-42, Thr-206, and Arg-211. In NgBR, Arg-290 is part of the conserved C-terminal -RXG- motif. (B) The side chain of Thr-206 is involved in hydrogen bonding (shown as dashed lines) with the backbone amide and carbonyl of Asp-34. The carbon atoms are colored deep teal, nitrogen atoms colored blue and oxygen atoms colored red. IPP1 colored red, IPP2 colored green and the Mg^2+^ is shown as a gray sphere.

**Supplementary Table 1.** Crystallographic statistics. The NgBR/DHDDS heterodimer was crystallized in space group R32.

| Data Collection |  |
| --- | --- |
| Wavelength, Å | 0.979 |
| Cell Dimensions, Å | a=b=185.7, c=113.4 |
| <sup>a</sup>Resolution (Å) | 40-2.5 (2.59-2.50) |
| Redundancy | 12.8 (11.2) |
| Completeness (%) | 99.2 (94.3) |
| <I/σ> | 7.5 (0.7) |
| <sup>a,b</sup>R<sub>merge</sub> | 0.108 (0.877) |
| Refinement |  |
| Unique reflections | 24,448 |
| Number of Atoms | 4,055 |
| Protein | 3,937 |
| Ligand | 29 |
| Solvent | 89 |
| <sup>c</sup>R<sub>work</sub>/R<sub>free</sub> | 0.201/0.289 |
| Mean B-value (Å<sup>2</sup>) | 69.8 |
| r.m.s. deviations |  |
| Bond lengths (Å) | 0.007 |
| Bond angles (°) | 1.760 |
| PDB accession code | 6W2L |

<sup>a</sup>Highest resolution shell is shown in parentheses.
<sup>c</sup>R<sub>work</sub> = $\sum \left| F_o - F_c \right| / \sum F_o$ . R<sub>free</sub> is the cross-validation R factor for the test set of reflections (5% of the total) omitted in model refinement.

**Supplementary Table 2.** Plasmids For *S.cerevisiae* expression system

| Source | *S.cerevisiae* plasmids | Description |
| --- | --- | --- |
| Park et al 2014 | pKG-GW1 | *LEU2* marker, 2 μ origin of replication |
| Park et al 2014 | pKG-GW2 | *MET15* marker, 2μ origin of replication |
| Park et al 2014 | pKG-GW2-NgBR | Derivative pKG-GW2, express NgBR gene |
| Park et al 2014 | pKG-GW1-hCIT | Derivative of pKG-GW1, express DHDDS gene |
| This study | pKG-GW1- DHDDS-256 | Derivative pKG-GW1-hCIT, express truncated version of 1-256 DHDDS |
| This study | pKG-GW1- DHDDS-289 | Derivative pKG-GW1-hCIT, express truncated version of 1-289 DHDDS |
| This study | pKG-GW1- DHDDS<sup>3A</sup> | Derivative pKG-GW1-hCIT, express mutated DHDDS<sup>R305A,F313A,F317A</sup> |
| This study | pKG-GW1- DHDDS<sup>DEKE</sup> | Derivative pKG-GW1-hCIT, express mutated DHDDS<sup>DEKE</sup> |

**Supplementary Table 3.** Plasmids for mammalian expression system

| Source | Mammalian expression vectors | Description |
| --- | --- | --- |
| Invitrogen | pcDNA3.1 | Mammalian expression vector with Cytomegalovirus (CMV) enhancer-promoter |
| Park et al 2014 | pcDNA3.1-FLAG-DHDDS | Derivative of pcDNA3.1 expressing N-terminally FLAG tagged DHDDS |
| Park et al 2014 | pcDNA3.1-NgBR-HA | Derivative of pcDNA3.1 expressing C-terminally HA tagged NgBR |
| This study | pcDNA3.1-NgBR-HA<sup>Q180A</sup> | Derivative pcDNA3.1-NgBR-HA expressing NgBR<sup>Q180A</sup> |
| This study | pcDNA3.1-NgBR-HA<sup>G196A</sup> | Derivative pcDNA3.1-NgBR-HA expressing NgBR<sup>G196A</sup> |
| This study | pcDNA3.1-NgBR-HA<sup>K197A</sup> | Derivative pcDNA3.1-NgBR-HA expressing NgBR<sup>K197A</sup> |
| This study | pcDNA3.1-NgBR-HA<sup>I200A</sup> | Derivative pcDNA3.1-NgBR-HA expressing NgBR<sup>I200A</sup> |
| This study | pcDNA3.1-NgBR-HA<sup>L226A</sup> | Derivative pcDNA3.1-NgBR-HA expressing NgBR<sup>L226A</sup> |
| This study | pcDNA3.1-NgBR-HA<sup>L230A</sup> | Derivative pcDNA3.1-NgBR-HA expressing NgBR<sup>L230A</sup> |
| This study | pcDNA3.1-NgBR-HA<sup>P238A</sup> | Derivative pcDNA3.1-NgBR-HA expressing NgBR<sup>P238A</sup> |
| This study | pcDNA3.1-NgBR-HA<sup>K243A</sup> | Derivative pcDNA3.1-NgBR-HA expressing NgBR<sup>K243A</sup> |
| This study | pcDNA3.1-NgBR-HA<sup>D248A</sup> | Derivative pcDNA3.1-NgBR-HA expressing NgBR<sup>D248A</sup> |
| This study | pcDNA3.1-NgBR-HA<sup>G252A</sup> | Derivative pcDNA3.1-NgBR-HA expressing NgBR<sup>G252A</sup> |
| This study | pcDNA3.1-NgBR-HA<sup>F253A</sup> | Derivative pcDNA3.1-NgBR-HA expressing NgBR<sup>F253A</sup> |
| This study | pcDNA3.1-NgBR-HA<sup>P255A</sup> | Derivative pcDNA3.1-NgBR-HA expressing NgBR<sup>P255A</sup> |
| This study | pcDNA3.1-NgBR-HA<sup>L260A</sup> | Derivative pcDNA3.1-NgBR-HA expressing NgBR<sup>L260A</sup> |
| This study | pcDNA3.1-NgBR-HA<sup>I263A</sup> | Derivative pcDNA3.1-NgBR-HA expressing NgBR<sup>I263A</sup> |
| This study | pcDNA3.1-NgBR-HA<sup>G292A</sup> | Derivative pcDNA3.1-NgBR-HA expressing NgBR<sup>G292A</sup> |

**Supplementary Table 4.** Plasmids for *E.coli* expression system

| Source | *E. coli* expression vectors | Description |
| --- | --- | --- |
| Addgene | pSUMO1 | Bacterial Expression of human 6His-tagged SUMO1 (#52258) |
| Novagen | pRSF-DUET1 | Bacterial dual expression vector, kan<sup>R</sup>, |
| This study | pRSF-DUET1-DHDDS/Sumo-NgBR | Derivative of pRSF-DUET1, express N-terminally taged 6-His Sumo-NgBR and untagged DHDDS |
| This study | pRSF-DUET1-DHDDS/Sumo-NgBR<sup>G91C</sup> | Derivative of pRSF-DUET1-DHDDS/Sumo-NgBR expressing NgBR<sup>G91C</sup> |
| This study | pRSF-DUET1-DHDDS<sup>W12A,F15A,I19A</sup>/Sumo-NgBR | Derivative of pRSF-DUET1-DHDDS/Sumo-NgBR expressing DHDDS<sup>W12A,F15A,I19A</sup> |

**Supplementary Table 5.**
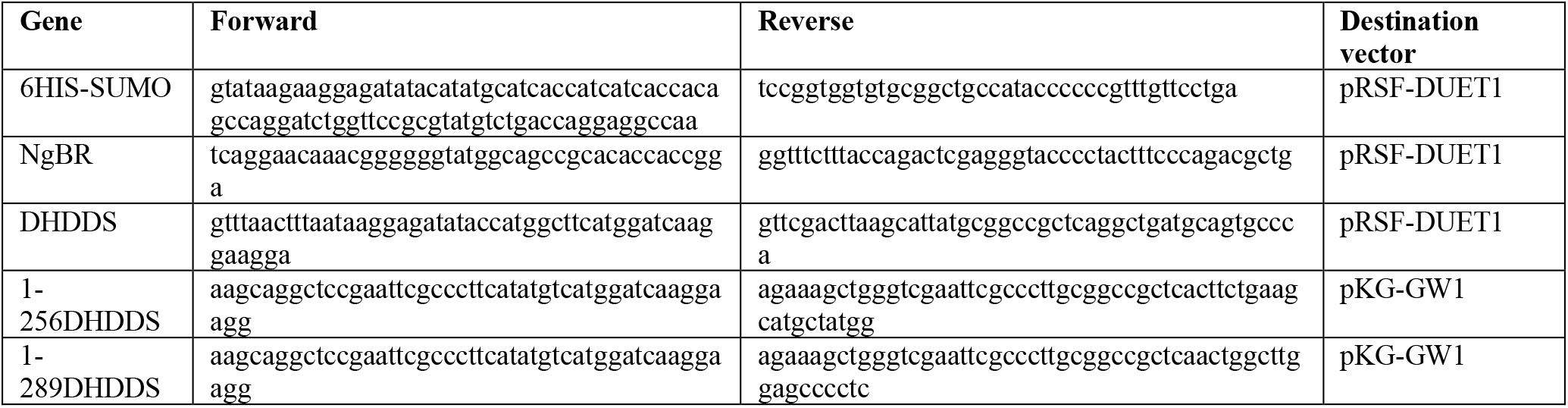
PCR primers used for cloning into expression vector

**Supplementary Table 6.**
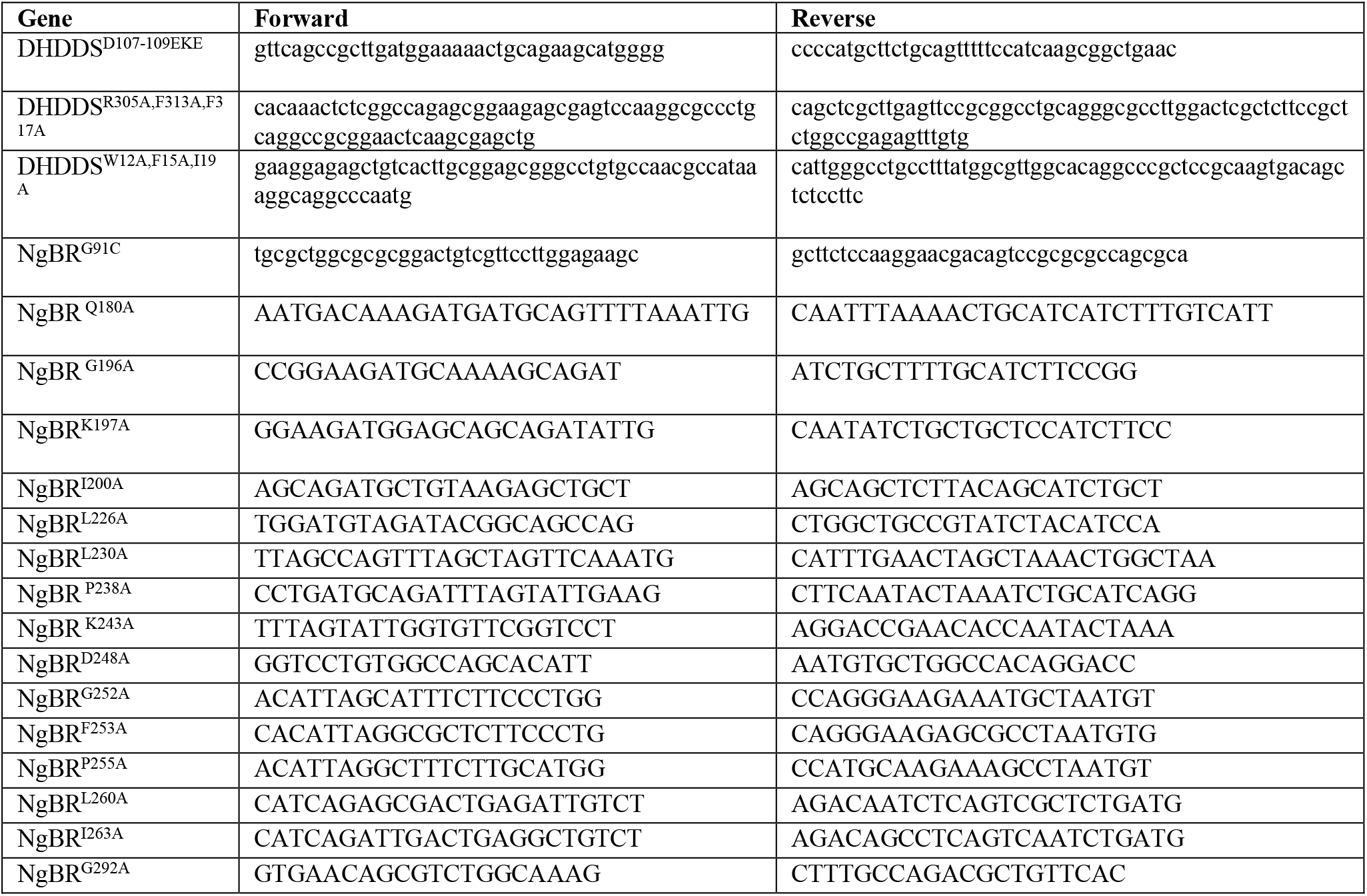
PCR primers used for site directed mutagenesis

